# Single cell mutant selection for metabolic engineering of actinomycetes

**DOI:** 10.1101/2022.01.13.476137

**Authors:** Amir Akhgari, Bikash Baral, Arina Koroleva, Vilja Siitonen, David P. Fewer, Charles E. Melançon, Jani Rahkila, Mikko Metsä-Ketelä

**Affiliations:** Department of Life Sciences, University of Turku, Turku FIN-20014, Finland; Department of Microbiology, University of Helsinki, Helsinki FIN-00014, Finland; Department of Chemistry and Chemical Biology, University of New Mexico, Albuquerque, New Mexico 87131-0001, USA; Instrument Centre, Faculty of Science and Engineering, Åbo Akademi University, Turku FIN-20500, Finland

**Author notes:** VTT Technical Research Centre of Finland Ltd., Tietotie 2, FIN-02044 Espoo, Finland. Applied Biomedical Science Institute, San Diego, CA 92127. Equal contribution (Amir Akhgari, Bikash Baral, Mikko Metsä-Ketelä). Correspondence and requests for materials should be addressed to Mikko Metsä-Ketelä.

**Keywords:** *Streptomyces*, *Amycolatopsis*, fluorescence-activated cell sorting, polyketide, protein production

## Abstract

Actinomycetes are important producers of pharmaceuticals and industrial enzymes. However, wild type strains require laborious development prior to industrial usage. Here we present a generally applicable reporter-guided metabolic engineering tool based on random mutagenesis, selective pressure, and single-cell sorting. We developed fluorescence-activated cell sorting (FACS) methodology capable of reproducibly identifying high-performing individual cells from a mutant population directly from liquid cultures. Genome-mining based drug discovery is a promising source of bioactive compounds, which is complicated by the observation that target metabolic pathways may be silent under laboratory conditions. We demonstrate our technology for drug discovery by activating a silent mutaxanthene metabolic pathway in *Amycolatopsis*. We apply the method for industrial strain development and increase mutaxanthene yields 9-fold to 99 mg l^−1^ in a second round of mutant selection. Actinomycetes are an important source of catabolic enzymes, where product yields determine industrial viability. We demonstrate 5-fold yield improvement with an industrial cholesterol oxidase ChoD producer *Streptomyces lavendulae* to 20.4 U g^−1^ in three rounds. Strain development is traditionally followed by production medium optimization, which is a time-consuming multi-parameter problem that may require hard to source ingredients. Ultra-high throughput screening allowed us to circumvent medium optimization and we identified high ChoD yield production strains directly from mutant libraries grown under preset culture conditions. In summary, the ability to screen tens of millions of mutants in a single cell format offers broad applicability for metabolic engineering of actinomycetes for activation of silent metabolic pathways and to increase yields of proteins and natural products.

## Introduction

Actinomycetes are filamentous Gram-positive bacteria that are widely utilized in diverse industrial biotechnology applications. Approximately two thirds of antibiotics and one third of anti-cancer agents in clinical use are natural products produced by actinomycetes or their semi-synthetic derivatives (1). Microbial natural products are widely used as immunosuppressants (2), antiparasitic (3) and antifungal agents (4). However, improved biologically active compounds are urgently needed to combat the emergence of antibiotic resistance in microbial pathogens (5) and to alleviate the side-effects commonly associated with cancer chemotherapy agents (6). Actinomycetes are likely to remain a critical source of new drug candidates (7), since genome sequencing projects have identified a tremendous number of unstudied biosynthetic gene clusters (BGCs) that could encode novel secondary metabolites (8). Methods for the bioinformatic identification of BGCs have become increasingly sophisticated (9), but isolation of the associated natural products has remained highly challenging. Genome mining driven drug discovery is complicated, because the majority of BGCs are silent under laboratory monoculture conditions. Many pathways require specific environmental signals for activation and are governed by complex regulatory cascades (10). Several techniques for activation of BGCs have been developed, but to date no single widely applicable methodology for targeted activation of metabolic pathways has emerged (11).

Actinomycetes play an important role in soil ecology, where they have adapted to decompose complex organic plant, anthropod and crustacean polymers (12). The proteins involved in these processes provide a significant fraction of commercially available enzymes used particularly in paper and pulp (e.g. xylanases and cellulases) and detergent (e.g. lipases and amylases) manufacturing (13). Numerous enzyme applications can also be found in the food and beverage (e.g. proteases and glucose oxidase) and textile (e.g. pectinases and peroxidases) industries (14). Many medical diagnostic laboratories also depend on enzyme use in bioassays (e.g. cholesterol oxidase) (15). Cholesterol oxidase ChoD is also utilized in the bioconversion of non-steroidal compounds, allylic alcohols and sterols, and in the chemoenzymatic synthesis of hormones and steroidal drug intermediates (15).

The manufacturing of both microbial natural products and proteins occurs in an enclosed bioreactor. Therefore product yields are key determinants for the commercial viability of the manufacturing process (16). Microbial strain development is an unceasing endeavor in pharmaceutical companies, where the yields of secondary metabolites have been increased from few tens of milligrams per liter in wild type strains to grams per liter in industrial production strains (17). Traditionally this decade-long process involves generation of millions of mutants by random mutagenesis followed by individual cultivation of each mutant strain to assess production profiles and yields (13). This last step is laborious and challenging to scale up even with high-throughput screening technologies. Once best producing mutants have been identified, the composition of medium ingredients needs to be optimized for high-yield production (13). This represents another time-consuming challenge, since medium optimization is a multi-parameter problem, where modification of one component influences the ideal concentrations of the other medium ingredients (16).

One attractive means of accelerating strain development has been to use reporter genes to focus on mutants with increased transcription of target genes (18–22). Increased yields of lovastatin (18) and natamycin (19) have been reported by linking transcription from the BGCs to the survival of the microbes under selection. A double reporter system was developed to reduce the number of false positive mutants to screen improved clavulanic acid producing mutants (20). Reporter guided mutant selection has also been utilized for the activation of silent BGCs with jadomycin (21), gaudimycin (21), streptothricin (22), geosmin (22) and strevertene (22) providing examples. Biosensors based on BGC situated repressor proteins have been used in the activation of coelimycin (23) pathway.

These studies have shown the value of reporter genes in strain development of actinomycetes. However, all depend on the detection of improved mutants on cultivation plates (16–19). This not only influences the throughput of the methodology, but also disassociates the screening conditions from the production environment in the bioreactor. Fluorescence-activated cell sorting (FACS) is an ideal method for sorting liquid cultures of heterogeneous mixtures of biological cells (24) and is routinely used in strain development of yeast (25) and selected bacteria (26). However, actinomycetes are filamentous bacteria that grow as branching mycelia, which hinders the use of FACS (27,28). A key problem is that the 70 μM nozzle size of the FACS instruments is typically orders of magnitude smaller than typical *Streptomyces* mycelia (e.g. 260-950 μM) (29,30). This limitation was circumvented by the use of protoplasts to pass through the instrument (27). However, the technique has ultimately not gained wide use, possibly because of the fragility of protoplasts and very low survival rates following cell sorting. Here we have developed facile methodology to allow reporter guided screening of actinomycetes by FACS. We demonstrate that single cell mutant selection (SCMS), which couples traditional mutagenesis to selection pressure and ultra-high throughput screening, can be used to activate silent BGCs, to increase the yields of proteins and secondary metabolites, and replace medium optimization.

## Results

### General workflow

A reporter construct is assembled in the first step by cloning the target promoter sequence in front of a double reporter plasmid pS-GK, constructed in the wide host range integrative pSET152 vector (31) (Fig. 1a and Supplementary Text). Two alternative versions with antibiotic resistance markers *kan* (32) or *hyg* (33) to impose selection using either kanamycin and hygromycin, respectively, are available. Importantly, the selection markers encode acyl transferase and kinase enzymes that inactivate their target antibiotics (34). This provides a link between the transcription level of the resistance gene and survival of the bacterial strain under elevated concentrations of the antibiotic. For the second reporter gene, we chose superfolder Green Fluorescent Protein (sfGFP) to allow screening by FACS. The reporter genes are insulated using two strong terminator sequences, a synthetic T4 kurz (35) and a natural terminator ECK120029600 (36). In the second step, classical mutagenesis is carried out to introduce genome-wide disturbances and to generate a mutant library (Fig. 1b). Positive mutants with increased transcription from the target promoter are enriched using selection (Fig. 1c) and subjected to ultra-high throughput screening by FACS to find best performing mutants (Fig. 1d). As a positive control for FACS, we devised plasmid pS-GK (Fig. 2a and Supplementary Fig. 1), where the reporter genes are cloned under the strong synthetic promoter SP44 (27). Pure cultures of mutants are initially ranked based on fluorescence (Fig. 1e), followed by quantification of target protein (Fig. 1f) or metabolite level (Fig. 1g).

**Fig. 1.**
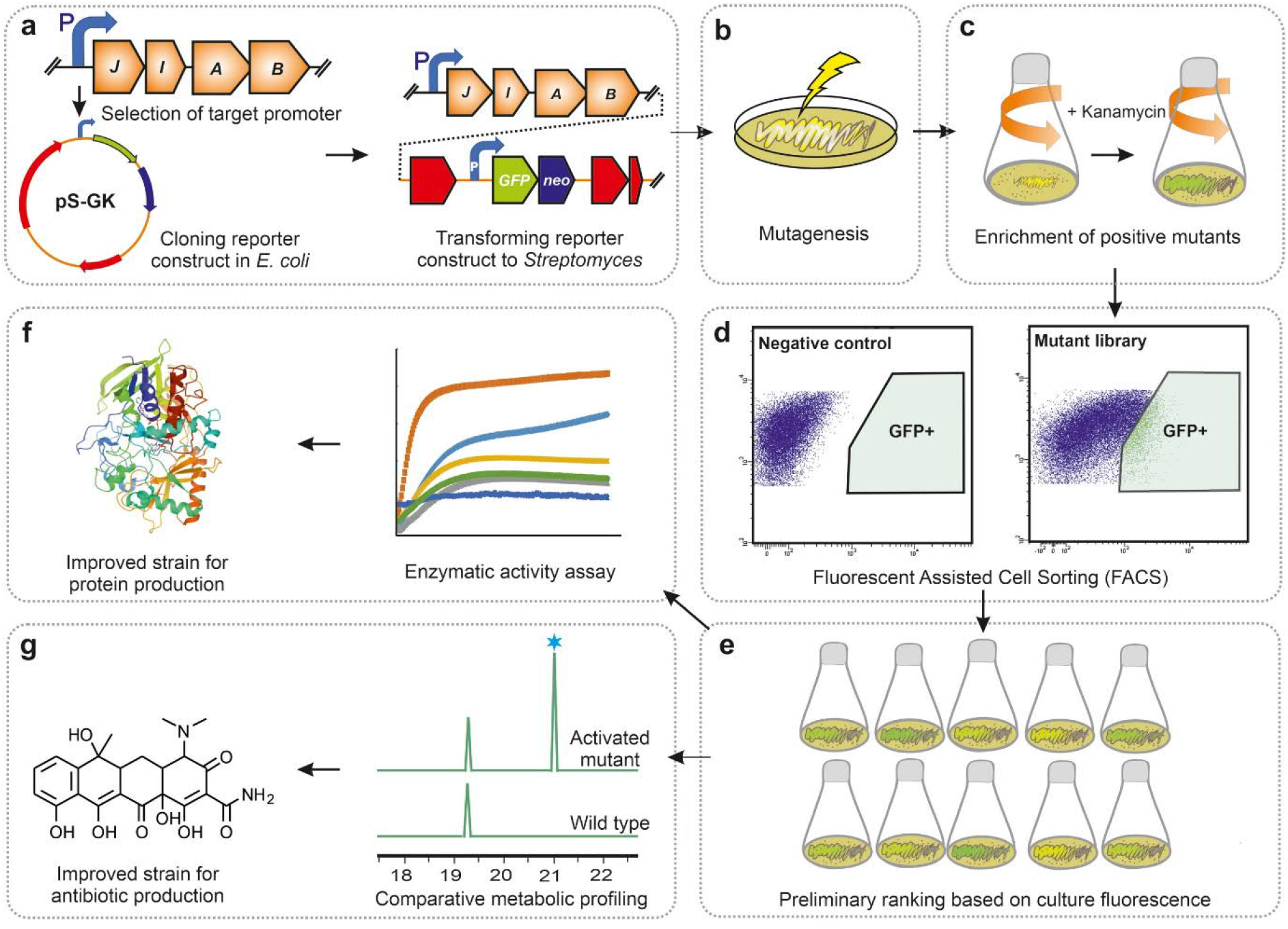
Single cell mutant selection platform for metabolic engineering. The strain improvement pipeline includes **a**, Construction of reporter plasmid by selecting a promoter region (P) from target gene(s), cloning the promoter in *E. coli* and conjugation of the plasmid into the production strain. **b**, Introduction of genome-wide disturbances by mutagenesis to generate overproducing strains. **c**, Transfer of the mutant library to a liquid culture and addition of antibiotics to impose selection to enrich positive mutants. **d**, Screening millions of mutants by FACS to find strains with highest fluorescence signal. Mutant libraries display a wide range of fluorescence values (x-axis) in comparison to a non-mutated negative control strain. Each dot represents a single cell. **e**, Preliminary analysis of pure cultures by fluorometer to rank mutants. **f**, Enzymatic activity assays to find best performing strain for protein overproduction or **g**, Comparative metabolic profiling to find improved strains for production of microbial natural products.

**Fig. 2.**
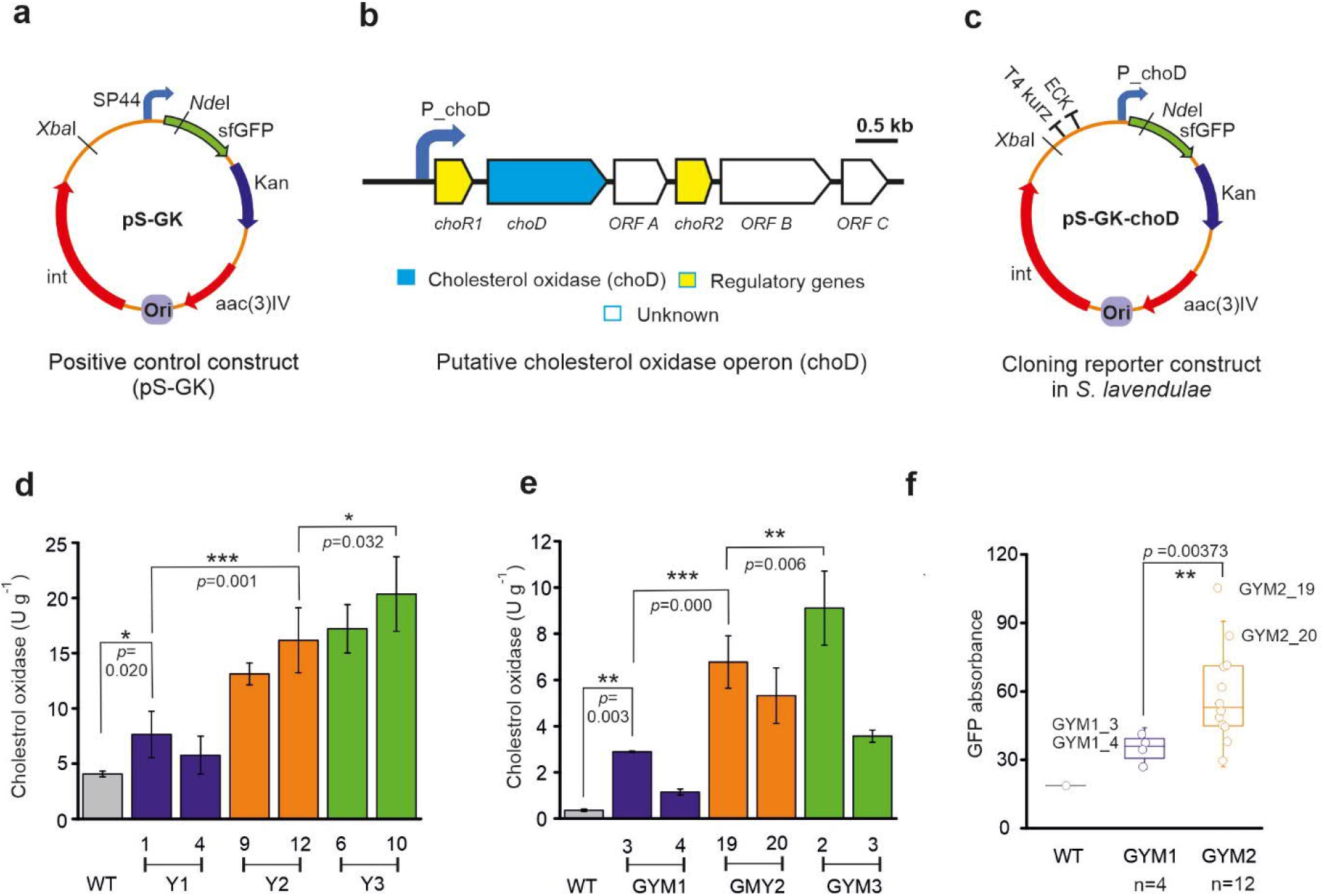
Improving ChoD protein production in *S. laevendulae* YAKB-15. **a**, Plasmid map for the positive control construct pS-GK containing the strong synthetic promoter SP44. Kanamycin resistance gene *kan* and *sfGFP* encoding superfolder Green Fluorescent Protein were used for selection and screening, respectively. Terminators T4 kurz^35^ and ECK120029600^36^ were used to insulate the promoter-probe construct. **b**, Structure of the *choD* operon and selection of the promoter region. **c**, The promoter for the *choD* operon was ordered as synthetic DNA and cloned as *Xba*I – *Nde*I fragment to generate the activation construct pS_GK_Chod. **d**, Cholesterol oxidase activity measurements from cultures grown in Y medium depicting increased yields of ChoD in three rounds of SCMS. Two mutants are shown for each round (Y1, first round; Y2, second round; Y3, third round) and the best performing mutant was selected for subsequent rounds of SCMS. **e**, Cholesterol oxidase activity measurements from cultures of two best mutants grown in GYM medium depicting increased yields of ChoD in three rounds of SCMS (GYM1, first round; GYM2, second round; GYM3, third round). **f**, Preselection of best stains after steak plating and fluorescence analysis in GYM medium. Lines depict the lineages of the strains in **d** and **e**. Significant differences were analyzed by one-way ANOVA with Tukey’s post hoc analysis, and *p*< 0.05 was considered statistically significant. \*\*\**p*< 0.001, \*\**p*< 0.01, \**p*< 0.05.

### Strain development for protein overproduction

First we focused on production of a cholesterol oxidase enzyme ChoD by an industrial strain *S. lavendulae* YAKB-15, which has been mutagenized for high yield production (12). The promoter sequence of the operon including *choD* (Fig. 2b) was cloned into the reporter construct in *Escherichia coli* and the resulting plasmid pS_GK_ChoD (Fig. 2c and Supplementary Fig. 2) was introduced to *S. lavendulae* YAKB-15. The exconjugants could withstand kanamycin concentrations up to 50 μg ml^−1^ in the production medium, reflecting the natural transcriptional level of *choD*. Next we generated a mutant library of *S. lavendulae* YAKB-15/pS_GK_ChoD by treatment of spores with ethyl methanesulfonate. The mutant library was used to inoculate parallel cultivations in Y production medium (15) supplemented with varying concentrations (50–400 μg ml^−1^) of kanamycin, with the objective of reducing the library size by enriching best performing mutants and killing off undesired low or non-producing mutants. Bacterial growth could still be observed in cultures with 400 μg ml^−1^ kanamycin, which demonstrated an 8.0-fold increase in the tolerance of the mutant library towards the antibiotic.

In order to utilize FACS, we hypothesized that fragmentation of the mycelium by sonication might be sufficient to allow single cells to pass through the nozzles of the instruments. Coupled with sample filtration and careful instrument gating (Supplementary Figs. 3-4), this simple method allowed us to analyze actinomycetes by FACS. We screened approximately 2 × 10^7^ cells from the enriched mutant library to find individual mutants with the highest sfGFP fluorescence. Positive mutants expressing sfGFP were harvested and analyzed for cholesterol oxidase activity in flask cultures in Y production medium (Fig. 2d). The mutant *S. lavendulae* Y1(1) displayed 1.8-fold increase in yield of ChoD (7.7 U g^−1^) in comparison to *S. lavendulae* YAKB-15 (4.1 U g^−1^).

Next we performed two more rounds of SCMS using the best performing mutant from the previous round as starting material. Single cells derived from second and third round mutagenesis and selection were resistant to 800 μg ml^−1^ and 900 μg ml^−1^ kanamycin, respectively. In total, three rounds of SCMS resulted in 5.0-fold yield enhancement in mutant *S. lavendulae* Y3(10) (20.4 U g^−1^) in comparison to *S. lavendulae* YAKB-15.

### Replacing the need for medium optimization

The production of cholesterol oxidase ChoD is tightly regulated in *S. lavendulae* YAKB-15 and is linked to the presence of whole yeast cells in the Y culture medium (12). Establishing optimal culture conditions is a necessity for industrial production of natural products and proteins by actinomycetes, since production levels fluctuate greatly in response to changing conditions (16). Since this considerably complicates media optimization, we concluded that the ability to take advantage of selection in strain development could be used to change the paradigm: instead of finding optimal culture conditions, SCMS could be used to find individual mutants that have a tendency for high yield production under preset culture conditions. Selection would allow identification of mutants that have adapted for high-yield production under conditions conforming to industrial conventions.

We selected yeast extract-based GYM medium, where the production yields of ChoD (0.4 U g^−1^) are low in *S. lavendulae* YAKB-15. The three rounds of single cell mutant selection led to continuously improved product titers (Fig. 2e) and generation of a mutant strain *S. lavendulae* GYM3(2), where the production of ChoD was 2.2-fold higher (9.1 U g^−1^) than from *S. lavendulae* YAKB-15 under optimized culture conditions (4.1 U g^−1^). The mutant libraries were found to be resistant to 200 μg ml^−1^, 400 μg ml^−1^ and 600 μg ml^−1^ kanamycin in the first, second and third round of mutation, respectively, demonstrating the importance of the enrichment steps.

During the strain development we noted fluctuations in ChoD production levels in strains acquired from *S. lavendulae* GYM1 and GYM2 mutant libraries, which we surmised might be due to acquisition of duplets instead of single cells by FACS. Product yields were stabilized through introduction of a preliminary ranking step based on overall fluorescence (Fig. 2f) of pure cultures obtained by streak plating. Importantly, higher fluorescence values (e.g. GYM1_3 and GYM2_19) correlated well with higher ChoD yields (Fig. 2e and 2f).

### Activation of a silent metabolic pathway

We used the software package *Dynamite* (37) to identify all putative type II polyketide BGCs in the NCBI databank. The resulting ~500 unstudied BGCs were scored and prioritized for predicted chemical novelty. Among these, the target BGC in *Amycolatopsis orientalis* NRRL F3213 was assessed to be transcriptionally silent based on the absence of typical type II polyketide UV/Vis spectral signatures in GYM medium extracts. The BGC (Fig. 3a and Supplementary Table 1) was predicted to encode a tetracycline-type compound based on ketosynthase (KS) α and β active site motifs and the complement of cyclase genes including a bifunctional aromatase-cyclase and *oxyN*-like cyclase genes (38). However, the presence of a priming ketosynthase III (KSIII) from an uncharacterized clade along with the absence of an *oxyD*-like amidotransferase indicated a new non-acetate primed tetracycline core structure, while the presence of a high number of oxidative and reductive enzymes suggesting extensive tailoring of the polyketide.

**Fig. 3.**
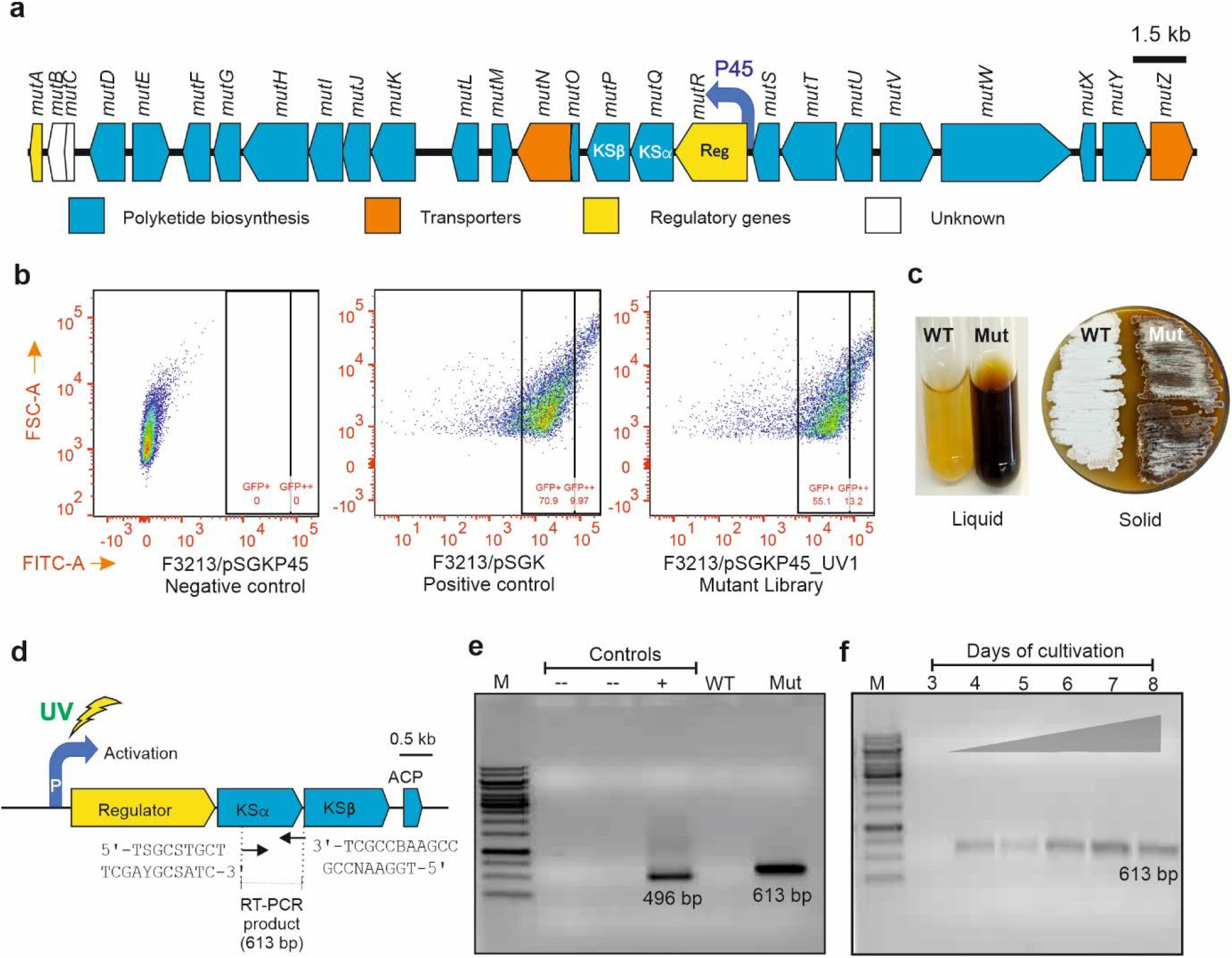
Activation of a mutaxanthene BGC in *A. orientalis* NRRL F3213. **a**, Organization of the aromatic type II polyketide BGC detected in *A. orientalis* NRRL F3213. The promoter targeted in SCMS is highlighted. **b**, FACS analysis of an enriched mutant library of *A. orientalis* NRRL F3213/pSGKP45 demonstrated enhanced sfGFP signal in a fraction of the mutants. Non-mutated *A. orientalis* NRRL F3213/pSGKP45 was used as a negative control, while *A. orientalis* NRRL F3213/pSGK, where the fluorescence signal is generated through expression of *sfGFP* from a strong constitutive synthetic promoter SP44, was used as a positive control. Each dot represents a single cell. **c**, Example of the phenotype of a GFP-positive mutant strain *A. orientalis* UV1(3) that produces pigmented metabolites in liquid cultures and on plates in contrast to the wild type strain. **d**, Schematic representation of the binding sites of primers used for detection of transcription from the core KS_α_ gene by RT-PCR. **e**, Confirmation of the cluster activation as detected by RT-PCR. Negative controls (lane 2 and 3) were reaction mixtures without reverse transcriptase enzyme and without RNA template, respectively. As a positive control, *in vitro*-transcribed human glyceraldehyde-3-phosphate dehydrogenase (GAPDH) control RNA was utilized (lane 4) that gave the product of size 496 bp. Detection of activated BGC on day 7 in *A. orientalis* UV1(3), where the expected 613 bp band is observed (lane 6). No products were observed in analysis of the wild type strain (lane 5). **f**, Time course analysis indicates that the BGC is activated on day 4 and transcription continues until day 8 in *A. orientalis* UV1(3).

We proceeded to clone the promoter region of an operon controlling the expression of the SARP-family regulatory gene (39) and the essential, translationally coupled, KS_α_ and KS_β_ responsible for synthesis of the polyketide scaffold, to generate the activation construct pSGKP45 (Fig. 3a and Supplementary Fig. 5) in *E. coli*. Introduction of the plasmid to *A. orientalis* NRRL F3213 did not induce formation of a sfGFP signal, reinforcing our hypothesis that the BGC was not transcribed under laboratory conditions. In order to activate the metabolic pathway, we proceeded to carry out mutagenesis of spores by UV radiation, enrichment under kanamycin selection (200 μg ml^−1^) and cell sorting by FACS (Fig. 3b). Single cells with positive sfGFP signal were harvested and grown in liquid cultures and on plates (Fig. 3c), which revealed phenotypically distinct mutants producing dark pigmented metabolites. We performed transcriptional analysis the KS_α_ gene by RT-PCR in order to validate that the targeted pathway had been activated in the mutant (Fig. 3d-f). The experiments confirmed that the BGC was active in the *A. orientalis* UV1(3) mutant, but not in the wild type (Fig. 3e). Time-course analysis indicated that the activation occurred on day four and the pathway remained active until end of the cultivation period on day eight (Fig. 3f).

Comparative metabolic analysis of culture extract by HPLC-UV/Vis (Fig. 4a) revealed that *A. orientalis* UV1(3) produced two main secondary metabolites **1** and **2** (Fig. 4b) not synthesized by the parental wild type strain. Furthermore, we noted that **1** was converted to **3** (Fig. 4b) during extractions in the presence of ammonium acetate. The mutant strain was cultivated in large-scale to obtain sufficient material for structure elucidation by NMR (Supplementary Tables 2-6) and HR-MS (Supplementary Fig. S6). The combined 1H and 13C, HSQCED, HMBC and 1,1-ADEQUATE NMR data confirmed that **1**, **2** and **3** were known polyketide natural products mutaxanthenes A, B and D, respectively (40). Conversion of **1** to its eneamine congener **3** in pH ~6.6 ammonium acetate buffer has been noted previously (30).

**Fig. 4.**
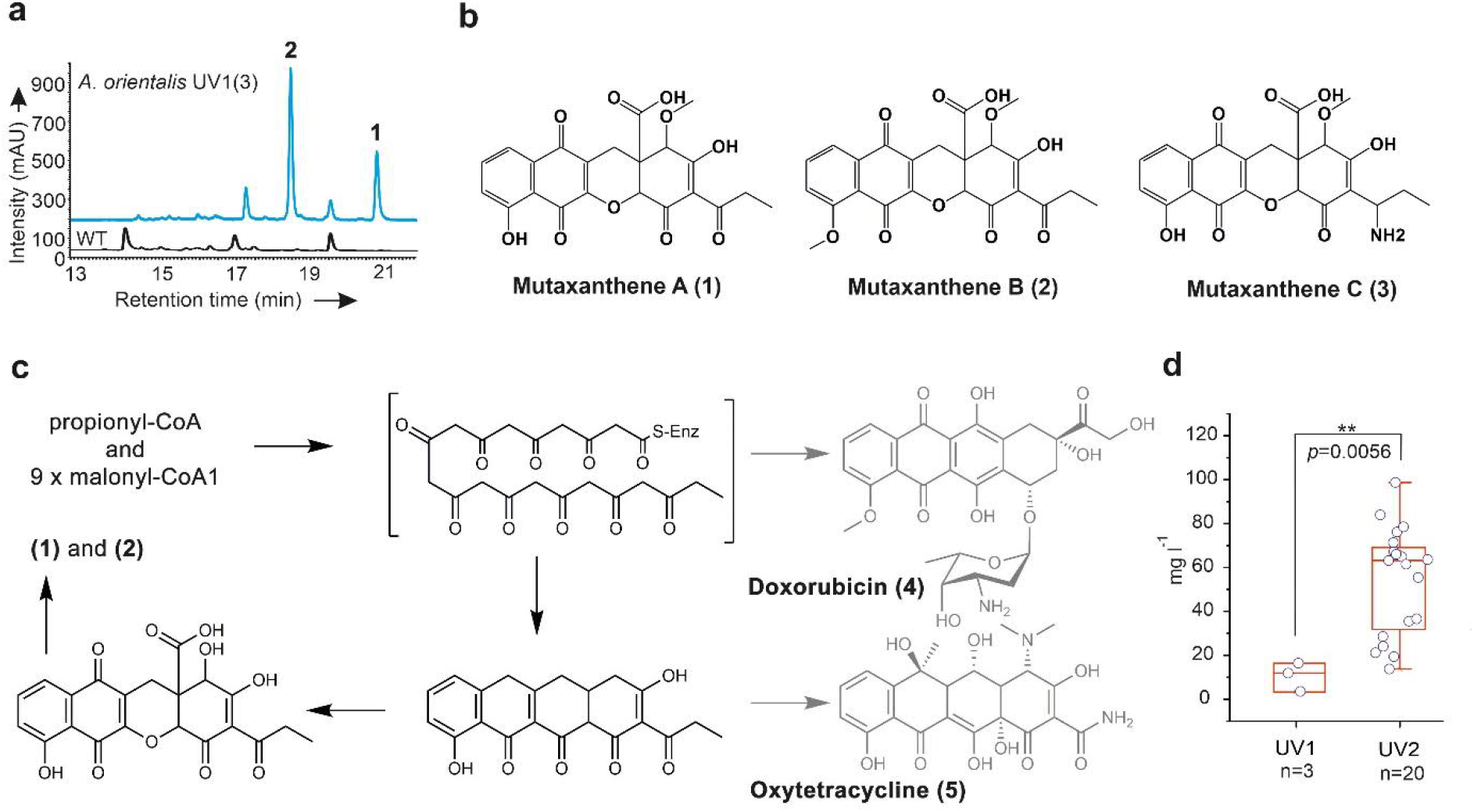
Structure elucidation, biosynthesis and yield improvement of mutaxanthenes. **a**, Analysis of culture extracts by HPLC-UV/Vis indicates the production of two additional metabolites **1** and **2** in *A. orientalis* UV1(3) in comparison to the wild type. The chromatogram traces were recorded at 256 nm. **b**, Chemical structures of mutaxanthenes A (**1**), B (**2**) and D (**3**) obtained from in *A. orientalis* UV1(3). **c**, The polyketide scaffolds of **1** and **2** are derived from propionyl-CoA started unit and nine malonyl-CoA extender units as in the biosynthesis of **4**. The set of cyclases encoded in the BBC that determine polyketide folding are related to the biosynthesis of **5**. A unique set of redox tailoring enzymes are utilized for multiple skeletal rearrangements that results in formation of **1** and **2**. **d**, Analysis of production profiles of 20 second round GFP-positive mutants demonstrates increased yields of **1** and **2**. Peak areas for the two metabolites obtained by HPLC were combined to reflect the total carbon flux to the activated pathway.

### Biosynthesis of mutaxanthenes

Analysis of the *mut* BGC provided further confirmation that the activated pathway is responsible for mutaxanthene biosynthesis. A mutaxanthene biosynthetic pathway has previously been reported in *Nocardiopsis* sp. FU40, but the nucleotide sequence was not deposited in public sequence repositories (40). Comparison of sequences obtained from the authors (Brian O. Bachmann, personal communication) revealed that the minimal polyketide synthases (KS_α_, KS_β_ and acyl carrier protein) of the two BGCs shared high 96.5% nucleotide sequence identity. As predicted, the biosynthesis is initiated by condensation of a non-acetate propionyl-CoA starter unit and nine malonyl-CoA extender units to produce an unreduced decaketide (Fig. 4c), similarly to the metabolic pathway of the anticancer agent doxorubicin (41) (**4**). A conserved set of cyclases fold the polyketide in a manner similar to oxytetracycline (**5**) biosynthesis (38). Deviations from the oxytetracycline paradigm are due to three additional genes that encode FAD-dependent proteins (Supplementary Table 1). These redox enzymes are likely to be responsible for the multiple skeletal rearrangements required for biosynthesis of mutaxanthenes.

### Improving yields of mutaxanthenes

The structures of **1** and **3** differ only in regards to 7-O-methylation and the combined average product titer of mutaxanthenes was calculated to be on average 11 mg l^−1^ in the three first round mutant strains. In order to improve production, we performed a second round of SCMS and acquired 40 pure culture mutants to estimate the robustness of the methodology. Twenty mutants were selected by ranking them based on culture fluorescence and the metabolic profiles of these strains were estimated by HPLC-UV/Vis. The second round strains displayed significantly increased yields with an average production of 55 mg l^−1^ (Fig. 4d). The best performing mutant *A. orientalis* UV2(38) demonstrated 9.0-fold increased yields of 99 mg l^−1^ in comparison to the parental first round mutant *A. orientalis* UV1(3).

## Discussion

Biomanufacturing provides an important means for production of biomolecules for use in medicine and numerous industrial applications. The field is likely to significantly grow in the future due to advances in synthetic biology and a shift to green chemistry (42). The commercial feasibility and success of these processes depend on product yields obtained from bioreactors. SCMS presents a generally applicable technology platform for industrial strain development of actinomycetes for production of pharmaceutical agents and industrial enzymes, where the efficiency of traditional strain development is increased by several orders of magnitude. Genome-wide mutagenesis is a proven method for increasing product yields (13), but numerous cumulative mutations with significant variation between production strains (43,44) are typically required for generation of an overproducing phenotype. SCMS can probe large mutant libraries directly from liquid production medium to identify best candidate strains. The use of selection in the removal of unwanted non- and low producing strains appears to be highly efficient, as demonstrated by the existence of very few individual mutant cells that are sfGFP negative in our mutagenesis libraries (Fig. 3b). Screening by FACS allows detection of the most promising strains from tens of millions of mutants. The combination of these two steps can be used to rapidly focus strain development efforts on the most promising fraction of the mutant library.

Actinomycetes are widely used for production of industrial enzymes due to good yields and protein solubility issues encountered in heterologous hosts (45). We show that SCMS can be used to efficiently increase product titer of the cholesterol oxidase ChoD. The 5.0-fold yield enhancement is noteworthy, since *S. lavendulae* YAKB-15 has been reported to have the highest activity of a cell-associated cholesterol oxidase (12). The combination of selection and ultra-high throughput screening may be sufficiently high for SCMS to replace medium optimization; we generated high-yield production strains in just three rounds of SCMS under preset culture conditions. The 22.8-fold yield enhancements obtained in GYM medium demonstrates the high dynamic range of the sfGFP reporter system. The strategy could be used to circumvent the need for exotic and hard to source medium ingredients required by industrial fermentation processes. One strain development cycle can be conducted in three-four weeks and iterative SCMS may provide a more cost effective approach for increasing production levels than conducting medium optimization experiments.

The discovery of numerous cryptic BGCs in actinomycetes has reinvigorated microbial natural product drug discovery pipelines (46). Computational genome mining is efficient in detection of potentially novel pathways (47), but robust methodologies for targeted activation of metabolic pathways have remained lacking. Cluster specific methodologies such refactoring (48), promoter exchange experiments (49) and manipulation of regulatory elements (50) require extensive investments and are not easily scaled to handle multiple samples. Ideally, the technology should be able to probe entire genomes to induce changes at global regulatory levels, but also target specific BGCs identified by genome mining. Currently existing methodologies include variations of small molecule elicitors (51,52), biosensors (53) and other reporter guided (54) systems. The ability to analyze large mutant libraries in SCMS will provide significant benefits over previous technologies, since activation of a metabolic pathway is likely to be a rare event resulting from a combination of several mutations. Importantly, the SCMS protocol (Fig. 1) allows high-throughput work with multiple BGCs in parallel. Activation of the silent mutaxanthene pathway and yield enhancement via iterative rounds of SCMS to a yield of 99 mg l^−1^ demonstrates that SCMS can facilitate identification of metabolites from culture extracts and provide sufficient material for structure elucidation and bioactivity assays. In conclusion, SCMS is a new tool that offers a wide range of metabolic engineering applications to accelerate drug discovery and industrial production of proteins and natural products.

## Methods

### Reagents

All reagents were purchased from Sigma-Aldrich unless otherwise stated. All organic reagents used for HPLC and HR-MS were high-performance liquid chromatography (HPLC) grade solvents.

### Strains and culture conditions

All plasmids were propagated (55) in *Escherichia coli* Top10 cultured in Luria-Bertani (LB) medium with apramycin (50 μg ml^−1^) or kanamycin (50 μg ml^−1^) at 37 °C. *E. coli* ET12567/pUZ8002 was used for conjugation (31) and grown in LB at 37 °C with appropriate antibiotics (25 μg ml^−1^ chloramphenicol, 50 μg ml^−1^ kanamycin, 50 μg ml^−1^ apramycin). To prepare spores, *Streptomyces lavendulae* YAKB-15 and *Amycolatopsis orientalis* NRRL F3213 were cultivated on mannitol-soya flour agar (31) plates at 30 °C.

### Construction of reporter plasmids

Codon-optimized oligonucleotide fragment of *sfgfp* with strong synthetic promoter (SP44) and ribosome binding site (SR41) (27) flanking *Xba*I/*Spe*I restriction sites (fragment I), kanamycin and hygromycin resistance genes with corresponding ribosome binding sites flanking *Spe*I/*Bam*HI restriction sites (fragment II) and two fragments consisting of two strong terminator and promoter region of *choD* or *mut* operon linked to *gfp* partial sequence (1-234 bp) flanked with *XbaI/Nde*I restriction sites (fragments III), were ordered as synthetic fragments (ThermoFisher Scientific). First pSET152 plasmid and fragment I were digested by *Xba*I*/Spe*I restriction enzymes and two purified fragments were assembled with T4 DNA ligase according to standard protocol (55). Then, fragment II was digested with *SpeI/Bam*HI and ligated to the plasmid backbone to create pS_GK construct (*gfp*+kanamycin resistance gene). Reporter plasmids were constructed by digestion of fragment III with *Xba*I/*Nde*I and ligation to similarly cut pS_GK. The constructs were transformed into *S. lavendulae* and *A. orientalis* via conjugations.

### Mutagenesis and selection

Chemical mutagenesis was carried out as described in Jones et al. (2017) (56) with slight modifications. Spores (*circa* 10^8^) of *S. lavendulae* harboring reporter probe were added to 1.5 ml of KPO_4_ (0.01 M; pH 7.0) and exposed to 200 μl ethyl methanesulfonate to achieve 99% killing rate while H_2_O was added to the control. The samples were vortexed for 30 seconds and incubated on shakers (300 rpm) at 30 °C for one hour, with inversions performed in 10-minute intervals. Then, the samples were centrifuged at 4000 rpm for 10 minutes at room temperature and subsequently the pellet was resuspended in one ml of freshly made and filter-sterilized 5% (w/v) sodium thiosulphate solution and then washed twice with 1 ml of H_2_O and subsequently the pellet was resuspended in 1 ml H_2_O. From each tube, a 10^−4^ dilution of the homogenate was made and 100 μl was plated onto MS plates and incubated for three days at 30 °C to estimate the killing rate. The rest of the homogenate was added to a 250 ml Erlenmeyer flask containing 25 ml of the antibiotic-free Y or GYM media and incubated on shaker at 300 rpm for 24 hours at 30 °C. The mutant library and control strains were sub-cultured (500 μl inoculation) in 25 ml fresh media supplemented with varying concentrations of kanamycin 50-900 μg ml^−1^ in three-day intervals.

### Quantitative Measurement of GFP Expression by Flow Cytometry

Mutant libraries of *S. lavendulae* and *A. orientalis* were subjected to FACS analysis to screen mutated single cells with the highest expression of GFP. Wild type strains and strains harboring strong *sfgfp* were used for FACS gating. Briefly, *Streptomyces* mycelia (3 day-old culture in 25 ml liquid media) were harvested (4 °C, 10,000*g*, 5 min) and then the pellet were washed with MQ water three times and subsequently was suspended in 25 ml PBS buffer. The pellet was then subjected to ultrasonication (amplitude 20, pulse 10 s: stop 10 s, 5 min, on ice) to generate mycelia fragments. Fragments were then filtered through 5 ml Falcon^®^ Polystyrene round bottom tubes with cell-strainer cap (Corning Science, Reynosa TAMP, Mexico). The fragmented mutated mycelia were analyzed by FACSAria Flow Cytometer with a 488-nm excitation laser and the FL1 (530/30 nm band-pass filter) detector. Each sample collected 50,000 events, and the data were acquired using BD FACSDiva™ 8.0.2 Software (BD Biosciences, CA) and analyzed by FlowJo™ v10.8 Software (BD Life Sciences). The parameters of the FACS setting were as follows: FSC-E00, SSC-650, FL1-400, FL2-400; threshold: FSC-50, SSC-400. The fluorescence of each sample was the geometric mean of all of the measured cells and was normalized to the corresponding FSC value, which indicates the size of the cells. Single cells were sorted in 96 well plate containing GYM media and were immediately plated on MS-agar plates. Several single colonies were streaked on MS media for three passages to guarantee obtaining pure culture.

### Cholesterol oxidase enzymatic assay

ChoD activity was measured spectrophotometrically as described previously (15). The stoichiometric formation of H_2_O_2_ during the oxidation reaction of cholesterol was monitored with ABTS at 405 nm. To determine the cell-bound ChoD, 500 μl of cultures were centrifuged at 15,000*g* for 10 min. The cell pellet was resuspended in extraction buffer (0.15% Tween 80 in 50 mM phosphate buffer solution) and mixed for 30 minutes at 4 °C. The suspension was centrifuged at 15,000*g* and ChoD activity was measured from the supernatant. The activity assay mixture contained 120 μl Triton X-100 (0.05%) in 50 mM sodium-potassium phosphate buffer (pH 7), 10 μl ABTS (9.1 mM in MQ H_2_O), 2.5 μl cholesterol in ethanol (1 mg ml^−1^), 1.5 μl horseradish peroxidase solution (150 U ml^−1^) and 20 μl of the extract preparation in a total volume of 154 μl. The spectrophotometric cholesterol activity assay was carried out in a 96-well plate. One unit of enzyme was defined as the amount of enzyme that forms 1 μmol of H_2_O_2_ per minute at pH 7.0 and 27 °C. All the samples including sorted cells, their corresponding control cultures and wild type strain were cultured in triplicates for 24 hours in corresponding media and then subcultured into fresh media for 72 hours (30 °C; 300 rpm).

### Production and purification of mutaxanthenes

To produce **1**-**3**, 50 μl of mutant spores’ stock (*A. orientalis*/pSGKP45_UV1(3)) were inoculated into 250 ml of TSB medium (2 flasks each with 250 ml) and grown for three days in a shaking incubator (30 °C; 300 rpm). The fully grown pre-culture was pooled together and 400 ml was transferred to 4 L of MS-broth and cultivated for 8 days (similar growth conditions) before being harvested. LXA resin was added to the cultures to bind produced secondary metabolites. After cultivation, compounds were extracted from LXA by first using 50% MeOH and then 90% MeOH. Both extracts were separately subjected to a liquid-liquid extraction by using CHCl_3_ first without adjusting the pH. The neutral CHCl_3_ was collected separately and the aqueous phase was further extracted twice with CHCl_3_ after adding 1% (v/v) acetic acid. The CHCl_3_ phases were dried and saved in −20 °C. The acidic CHCl_3_ was subjected to a silica column (diameter 5.6 cm and length 9 cm) equilibrated with following conditions: toluene:ethyl acetate:methanol:formic acid 50:50:15:3. The first colorful front was collected in 90 ml fractions and 10 ml of ammonium actetate (1M) was added. The first four fractions were extracted using 1M ammonium acetate, the compounds of interest were in the aqueous phase. Next the aqueous phase was acidified using formic acid and acetic acid and extracted with CHCl_3_. The CHCl_3_ phases were dried, dissolved in methanol and injected to preparative HPLC, LC-20 AP with a diode array detector SPD-M20A (Shimadzu) with a reverse-phase column (Phenomenex, Kinetex) using a gradient from 15% acetonitrile with 0.1% formic acid to 100% acetonitrile. The peaks of interest were collected, extracted with CHCl_3_ and dried under vacuum and with nitrogen gas. NMR samples were prepared from overnight desiccated samples in CDCl_3_ (**1**, **2**) or MeOD-*d4* (**3**).

### Analyses of compounds by HPLC, NMR and HR-MS

HPLC analyses were carried out with a SCL-10Avp HPLC with an SPD-M10Avp diode array detector (Shimadzu) using a reversed-phase column (Phenomenex, Kinetex, 2.6 μm, 4.6 × 100 mm) using a gradient from 15% acetonitrile with 0.1% formic acid to 100% acetonitrile. NMR analysis was performed with a 600 MHz Bruker AVANCE-III NMR-system equipped with a liquid nitrogen cooled Prodigy TCI (inverted CryoProbe) at 298 K. The experiments included 1D (^1^H, ^13^C) and 2D measurements (COSY, HMBC, HSQCED, NOESY, TOCSY and 1,1-ADEQUATE). Topspin (Bruker Biospin) was used for spectral analysis and accurate chemical shifts and coupling constants were extracted with ChemAdder (Spin Discoveries Ltd.). High resolution mass (MicroTOF-Q, Bruker Daltonics) was performed with direct injection of purified compounds.

### Transcriptional analysis by RT-PCR

To assess gene expression, *A. orientalis* NRRL F3213 wild type and an activated mutant (*A. orientalis*/pSGKP45_UV1(3)) were cultivated in 50 ml MS-liquid medium for 8 days. Sampling for RNA isolation was performed daily from 3d until 8d cultures. For this, 4 ml of the culture was harvested and gene expression was halted by addition of 444 μl stop solution (95% ethanol, 5% phenol). The mixture was incubated at room temperature for 15 min, followed by centrifugation (12,000*g*; 15 min). The pellets were flash frozen in liquid nitrogen for storage at −80 °C until RNA was extracted.

Total RNA was extracted using the RNeasy Mini Kit (Qiagen) according to the manufacturer’s instructions with minimal adjustments. Briefly, liquid nitrogen was employed to ground the cells to a powder. To the powder, 2 ml of phenol/CHCl_3_ /isoamyl alcohol (25:24:1) was added, followed by the addition of 1 ml TE (30mM Tris-Cl, 1mM EDTA, pH 8), and centrifuged (12,000*g*; 6 min). The aqueous layer was extracted for a second time with 700 μl CHCl_3_ and centrifuged (12,000*g*; 4 min). To the upper layer (*circa*. 500 μl), 400 μl RLT and 500 μl of ethanol were added successively. The mixture was loaded onto the column, rinsed with 350 μl RW1, and then 80 μl of DNase I digest was applied to the column, followed by incubation at room temperature for 30 min before being washed with RW1 (350 μl) and RPE (500 μl; 2 times). Eventually, the column was eluted with 80 μl RNase-free water. The extracted mRNA was checked for purity on a 0.8% agarose gel and stored at −80 °C until further use.

Synthesis of cDNA from the extracted mRNA was performed using the ThermoScientific RevertAid First Strand cDNA synthesis kit with modest adjustments. Briefly, the 12 μl reaction contained 4 μg of template RNA, 1 μl of random hexamer primer, and DEPC water. This reaction mixture was incubated at 65 °C for 5 min, before being cooled on ice. To this mixture, 5× reaction buffer (4 μl), RiboLock RNase inhibitor (20 U μl^−1^; 1 μl), 10 mM dNTP mix (2 μl) and reverse transcriptase (200 U μl^−1^; 1 μl) was added. The reaction mixture was incubated at 25 °C for 5 min, then at 42 °C for 65 min before being terminated by heating at 70 °C for 5 min.

RT-PCR amplification of the expressed gene targeting to the minimal polyketide synthase II gene regions was performed using the complementary DNA (cDNA) as a template and a pair of degenerate primers (5’-TSGCSTGCTTGGAYGCSATC-3’) (sense primer) and (5’-TGGAANCCGCCGAABCCGCT-3’) to amplify a product with the size of 613 bp (57). PCR product was analyzed by gel electrophoresis on a 0.8% agarose gel stained with SybrSafe and compared to a one kb Plus Ladder (Invitrogen). RT-PCR amplifications was performed for all the samplings harvested, viz., from 3d until 8d for both wild type and the mutants. The 50 μl of the reaction mixture for the RT-PCR amplification contained cDNA (1000× diluted): 1 μl; 10× GC buffer: 10 μl; 10 mM dNTP mix: 2.5 μl; DMSO: 2.5 μl; Primers (2.5 μl each for forward and reverse); Dream Taq DNA polymerase: 0.5 μl; and DEPC water: 28.5 μl. The thermocycling conditions comprise an initial longer denaturation phase (96 °C; 2 min). The cycle steps for 30 cycles were as follows: denaturation (96 °C; 1 min), annealing (58 °C; 2 min), and extension (73 °C; 1.5 min). The reaction was terminated with a longer final extension (73 °C; 8.5 min). RT-PCR was performed on a SureCyler 8800 (Agilent Technologies, Santa Clara, California, USA). Conditions for the control reactions are described in the Supporting Information Text.

## Accession numbers

The nucleotide sequences encoding the mutaxanthene were acquired from NCBI with the accession number JOIQ01000020.1. Nucleotide site for the ‘mut’ gene cluster was identified to be between 183729-216609. The biosynthetic gene cluster encoding mutaxanthene from A. orientalis NRRL F3213 was deposited in MIBiG (58) under the accession number BGC0002137.

## Statistical analyses

All error bars shown in the present work are standard deviation values of three biological replicates. The number of replicates is provided in the corresponding figure caption. One-way analysis of variance (ANOVA) was used to demonstrate differences in values.

## Acknowledgements

The financial support from the Jane and Aatos Erkko foundation to M.M.-K., the Finnish Cultural Foundation to A.A. and the Turku University Foundation to B.B. is acknowledged. The authors would like to thank Wubin Gao, Džesika Pozlevičiūtė and Sazia Rahman for assistance in experimental analyses. We thank Tiina Pessa-Morikawa for assistance in flow sorting performed at the HiLife Flow Cytometry Viikki Sorting Unit, University of Helsinki.

## Author Contributions

A.A. and B.B. conducted experiments, analyzed data and wrote the manuscript. A.K., V.S., C.M., D.P.F. and J.R. conducted experiments and analyzed data. M.M.-K. designed and conducted experiments, analyzed data, supervised the research and wrote the manuscript. All authors edited the manuscript.

