## Supporting information for "Single cell mutant selection for metabolic engineering of actinomycetes"

**Affiliations:**

± **Current address:** Applied Biomedical Science Institute, San Diego, CA 92127.

* **Equal contribution**

**Table of Contents:**

Supplementary Text……………………………………………………………………….2

Supplementary Figures 1-5………………………………………………………………..6

Supplementary Tables 1-6…………………………………………………………..…12

**Supplementary Text**

**Protocol for SCMS: Cloning Reporter Construct**

1. Design synthetic fragment III (Supplementary Figures S1, S2 and S5) comprising target promoter sequence surrounded by two terminators, RBS, part of the sf*gfp* gene and *Xba*I/*Nde*I restriction sites.
2. Digest plasmids harboring fragment III and pS-GK or pS-GH by *Xba*I/*Nde*I restriction enzymes. This digestion removes synthetic strong promoter and ribosome binding site and partial *gfp* sequence (the first 234 bp *gfp*) of pS-GK or pS-GH vector.
3. Clone fragments to yield a reporter plasmid with the target promoter sequence using apramycin (50 µg $ml^{-1}$) selection in *E. coli* TOP10.
4. Transform the plasmid to *E. coli* ET12567/pUZ8002 using apramycin (50 µg $ml^{-1}$), kanamycin (50 µg $ml^{-1}$) and chloramphenicol (25 µg $ml^{-1}$).
5. Conjugate reporter construct to target actinomycete strain and select exconjugants on MS agar containing nalidixic acid (25 µg $ml^{-1}$) and apramycin (50 µg $ml^{-1}$).

**Protocol for SCMS: Chemical Mutagenesis**

1. Grow *Streptomyces* strain harboring the reporter construct on MS-agar media supplemented with 50 µg $ml^{-1}$ apramycin for 3-5 days. In present work *S. lavendulae* transformed with a reporter construct harboring kanamycin resistance gene is presented as an example for chemical mutagenesis.
2. Prepare spore suspension and store in 20% sterile glycerol at -20 °C.
3. Transfer spores (*circa* $10^8$) of *Streptomyces* harboring the reporter probe to 1.5 ml of $KPO_4$ (0.01 M; pH 7.0).
4. Add 200 µl of ethyl methanesulfonate (EMS; Sigma-Aldrich, Inc., Product number M0880) to achieve more than 99% killing rate. The same amount of sterile MilliQ $H_2O$ was added to the control.
5. Vortex the samples for 30 seconds and incubate on shakers (300 rpm) at 30 °C for one hour, with inversions at every 10-minute interval.
6. Centrifuge samples at 4000 rpm for 10 minutes at room temperature and discard supernatant.
7. Resuspend the cell pellets in one ml of freshly made and filter-sterilized 5% w/v sodium thiosulphate ($Na_2S_2O_3 \cdot 5H_2O$; BDH, England) solution and then centrifuge (step 6).

8. Wash the pellets twice with one ml of MilliQ $H_2O$.
9. Subsequently, resuspend the pellets with one ml MilliQ $H_2O$.
10. Prepare a $10^{-4}$ dilution of each tube, plate 100 µl from the mutant library onto MS plates and incubate it for three days at 30 °C to estimate the killing rate.

**Protocol for SCMS: UV Mutagenesis**

1. Grow the *Amycolaptosis* strain harboring the reporter construct on MS-agar medium for 5 days supplemented with 50 µg $ml^{-1}$ apramycin. In this work, UV mutagenesis is demonstrated using *Amycolaptosis orientalis* NRRL F3213, which has been transformed with a reporter construct harboring the *gfp* + $kan^R$ genes.
2. Prepare spore suspension and store in 20% sterile glycerol at -20 °C.
3. In the laminar, transfer spores to sterile-plates beneath an UV-lamp at 10 cm. Use a magnetic stirrer to shake the spores, which allows aeration and UV light to reach all the spores.
4. Expose the spores to UV radiation (UV-C) for 15 min to attain more than 90% killing rate.

**Protocol for SCMS: Selection**

1. Transfer the mutant library to a 250 ml Erlenmeyer flask containing 25 ml of the antibiotic-free GYM medium and incubate on a shaker at 300 rpm for 24 hours at 30 °C. In case of UV mutagenesis, incubate the culture in dark incubators to prevent the mutation from reversing due to photolyase activity.
2. Make glycerol stock of the mutant library and store at -80 °C.
3. Determine the highest concentration of antibiotic, which can be tolerated by the mutant library and control cultures. Subculture 1.5 ml of control and mutant library samples in 25 ml of corresponding liquid medium (Y or GYM) supplemented with increasing amounts of antibiotic (e.g. 25, 50, 100, 200, 400 and 800 µg $ml^{-1}$ kanamycin) at three-day intervals.
4. The high kanamycin-tolerant mutant libraries and their corresponding non-mutated controls can be stored in 20% sterile glycerol at -80 °C.

**Protocol for SCMS: Flow Cytometry**

1. Prepare a 25 ml fresh liquid culture of mutant library, parental mutant (or wild type) strain from the previous round and a positive control strain harboring the pS-GK construct (expressing GFP under strong synthetic promoter) supplemented with antibiotics at the appropriate concentration. Grow them at 30 °C shaking at 300 rpm for 72 hours.
2. Harvest *Streptomyces* mycelia by centrifuging at 4 °C, 10,000*g* for 5 min.
3. Wash the pellet with 25 ml MilliQ water three times.
4. Subsequently resuspend the pellet in 25 ml of PBS buffer (137 mM NaCl, 2.7 mM KCl, 8 mM $Na_2HPO_4$, and 2 mM $KH_2PO_4$).
5. Subject the homogenized pellet to ultrasonication (amplitude 20, pulse 10 s: stop 10 s, 5 min, on ice) to generate mycelia fragments.
6. Filter the mycelial fragments suspension through 5 ml Falcon® Polystyrene round bottom tubes with cell-strainer cap (Corning Science, Reynosa TAMP, Mexico).
7. Sort individual cells in 96 well-plate GYM media (4 g $l^{-1}$ glucose, 4 g $l^{-1}$ yeast-extract, and 10 g $l^{-1}$ malt-extract).
8. Immediately plate sorted cells in MS-agar media (20 g $l^{-1}$ each of agar, mannitol and soya flour) supplemented with appropriate antibiotic concentrations and incubate at 30 °C for 3-7 days.
9. Streak plate to guarantee the axenic pure cultures. Let each passage grow for 5 days at 30 °C.
10. Make spore/mycelia suspension from single colonies and store in 20% sterile glycerol overnight at -20 °C before transferring to a -80 °C freezer for long-term storage.

**Control reactions for verifying first strand cDNA synthesis.** Positive and negative control reactions were used to validate the results of first strand cDNA synthesis. The experiment comprised two sets of negative controls: (i) reaction mixture without reverse transcriptase enzyme (to test genomic DNA contamination), and (ii) reaction mixture without template (to assess reagent contamination), were used independently. We used human glyceraldehyde-3-phosphate dehydrogenase (GAPDH) control RNA generated by *in vitro* transcription provided by the manufacturer (Thermo Scientific) as a positive control.

**Control first-strand cDNA synthesis reaction.** The 20 µl of control cDNA synthesis comprised control GAPDH RNA (2 µl), Random Hexamer Primer (1 µl); 5× reaction buffer (4 µl), RiboLock RNase Inhibitor (1 µl), 10 mM dNTP Mix (2 µl), Reverse transcriptase (1 µl), and nuclease-free water (9 µl). The mixed reaction was incubated at 25 °C for 5 min, then at 42 °C for 60 min, before being terminated by heating at 70 °C for 5 min. This reaction mixture was used diluted for 1000-folds and was used as a template for the PCR reaction.

**Control PCR reaction and thermocycling conditions.** The 50 µl of the reaction mixture for control PCR amplification contained cDNA (1000× diluted): 2 µl; 10× PCR buffer: 5 µl; 10 mM dNTP mix: 1 µl; $MgCl_2$ (25 mM; 3 µl); GADPH primers (1.5 µl each for forward and reverse); Taq DNA polymerase: 0.5 µl; and DEPC water: 35.5 µl. The thermocycling conditions comprised of longer denaturation phase (94 °C; 3 min), and the cycle steps for 35 cycles were as follows: denaturation (94 °C; 30s), annealing (58 °C; 30s) and extension (72 °C; 45). After the completion of the PCR reaction, 20 µl of the product was loaded on 0.8% agarose gel. The visibility of 496 bp PCR product confirmed the success of the RT-PCR reaction.

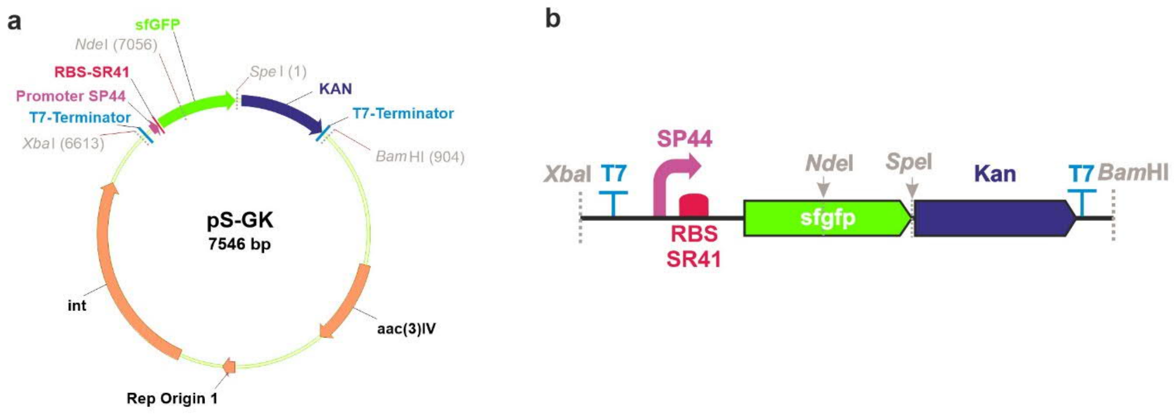

c

```
CTAGAGAGCTCGCTGAGTTGGCTGCTGCCACCGCTGAGCAATAACTAGCATAACCCCTTGGGGCCTCTAAACGGGTCTT
GAGGGGTTTTTTGCTGAAAGGAGGAACTATATCCGCGTGTTCACATTCGAACCGTCTCTGCTTTGACAACATGCTGTGC
GGTGTTGTAAAGTCTGGTGTAGGAGAATACGACAGTCTAAGTAAGGAGTAGGCTGATGCGCAAGGGCGAGGAGCTGTTC
ACCGGCGTCGTCCCGATCCTGGTCGAGCTGGACGGCGACGTCAACGGCCACAAGTTCTCCGTCCGCGGCGAGGGCGAGG
GCGACGCCACCAACGGCAAGCTGACCCTGAAGTTCATCTGCACCACCGGCAAGCTGCCGGTCCCGTGGCCGACCCTGGT
CACCACCCTGACCTACGGCGTCCAGTGCTTCGCCCGCTACCCGGACCATATGAAGCAGCACGACTTCTTCAAGTCCGCC
ATGCCGGAGGGCTACGTCCAGGAGCGCACCATCTCCTTCAAGGACGACGGCACCTACAAGACCCGCGCCGAGGTCAAGT
TCGAGGGCGACACCCTGGTCAACCGCATCGAGCTGAAGGGCATCGACTTCAAGGAGGACGGCAACATCCTGGGCCACAA
GCTGGAGTACAACTTCAACTCCCACAACGTCTACATCACCGCCGACAAGCAGAAGAACGGCATCAAGGCCAACTTCAAG
ATCCGCCACAACGTCGAGGACGGCTCCGTCCAGCTGGCCGACCACTACCAGCAGAACACCCCGATCGGCGACGGCCCGG
TCCTGCTGCCGGACAACCACTACCTGTCCACCCAGTCCGTCCTGTCCAAGGACCCGAACGAGAAGCGCGACCACATGGT
CCTGCTGGAGTTCGTCACCGCCGCCGGCATCACCCACGGCATGGACGAGCTGTACAAGTGATGAACTAGTGTAGCTGAC
AAGAGACAGGATGAGGATCGTTTCGCATGATTGAACAAGATGGATTGCACGCAGGTTCTCCGGCCGCTTGGGTGGAGAG
GCTATTCGGCTATGACTGGGCACAACAGACAATCGGCTGCTCTGATGCCGCCGTGTTCCGGCTGTCAGCGCAGGGGCGC
CCGGTTCTTTTTGTCAAGACCGACCTGTCCGGTGCCCTGAATGAACTGCAAGACGAGGCAGCGCGGCTATCGTGGCTGG
CCACGACGGGCGTTCCTTGCGCAGCTGTGCTCGACGTTGTCACTGAAGCGGGAAGGGACTGGCTGCTATTGGGCGAAGT
GCCGGGGCAGGATCTCCTGTCATCTCACCTTGCTCCTGCCGAGAAAGTATCCATCATGGCTGATGCAATGCGGCGGCTG
CATACGCTTGATCCGGCTACCTGCCCATTCGACCACCAAGCGAAACATCGCATCGAGCGAGCACGTACTCGGATGGAAG
CCGGTCTTGTCGATCAGGATGATCTGGACGAAGAGCATCAGGGGCTCGCGCCAGCCGAACTGTTCGCCAGGCTCAAGGC
GCACATACCCGACGGCGAGGATCTCGTCGTGACCCATGGCGATGCCTGCTTGCCGAATATCATGGTGGAAAATGGCCGC
TTTTCTGGATTCATCGACTGTGGCCGGCTGGGTGTGGCGGACCGCTATCAGGACATAGCGTTGGCTACCCGTGATATTG
CTGAAGAGCTTGGCGGCGAATGGGCTGACCGCTTCCTCGTGCTTTACGGTATCGCCGCTCCCGATTCGCAGCGCATCGC
CTTCTATCGCCTTCTTGACGAGTTCTTCTGAGCGGGACTCTGGGGTAGCATAACCCCTTGGGGCCTCTAAACGGGTCTT
GAGGGGTTTTTTGTTCGGGG
```

**Supplementary Figure 1. Organization of the positive control reporter construct pS-GK.** **a**, Plasmid map of pS-GK comprising terminators (T7), promoter (SP44), ribosome binding site (RBS-SR41) and double reporter genes (sf*gfp* and kanamycin) cloned to the pSET-152-based plasmid. **b**, Illustration of the construct along with the SP44 promoter cloned upstream of the reporter genes (sf*gfp* and kanamycin). Bacteriophage T7-terminator upstream of SP44 promoter insulates the transcription leakage. Promoter and its corresponding RBS drives the expression of sf*gfp* (super folder green fluorescent protein) and kanamycin resistance gene (*kan*). **c**, Depiction of nucleotide sequences of the region (*Xba*I/*Nde*I) highlighted in 'b' panel. Restriction enzymes (*Xba*I, *Nde*I, *Spe*I, and *Bam*HI) are highlighted with grey, T7-terminator with faint blue, promoter (SP44) underlined in purple, sf*gfp* with bright green, and resistance gene (*kan*) with dark blue.

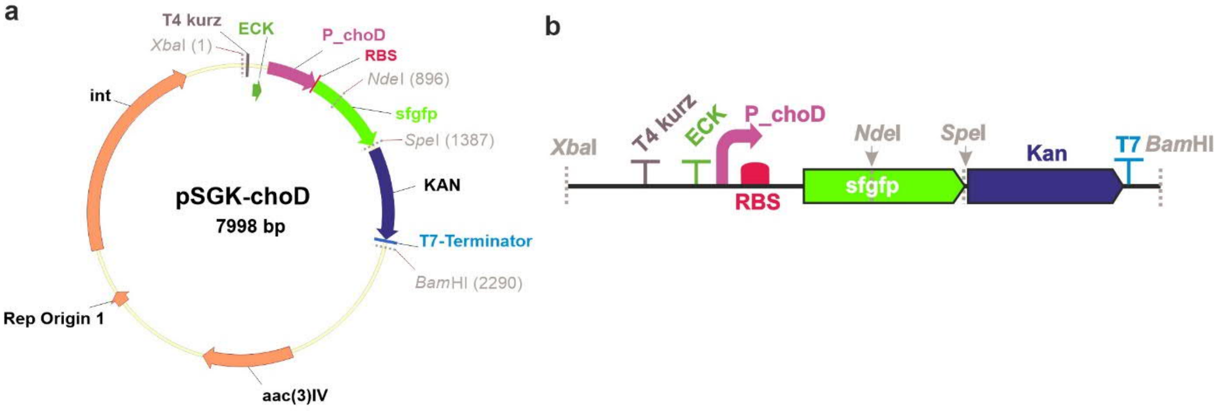

c

CTAGAGAGCTCGCTGAGTTGGCTGCTGCCACCGCTGAGCAATAACAAGCTTTATGCTTGTAAACCGTTTTGTGAAAAAA
TTTTTAAAATAAAAAAGGGGACCTCTAGGATTTTCAGCCAAAAAACTTAAGACCGCCGGTCTTGTCCACTACCTTGCAG
TAATGCGGTGGACAGGATCGGCGGTTTTCTTTTCTCTTCTCAACTGAAAGGAGGAACTATATCCGCGGGGCCCCAGCAC
CCTGAGGAGGGGCTCGGCCCGGCCGCTGGTCATGGGCGACGTGTTCACGGGAGGCGTGGACATGACGGTCCCCGATCTC
TTACGGGTACGGCTGGTTGACGGACGGCTTCGCGCAGGGGCTGTCGCAGGGCGTCCGCTACCGATCATGACATAGCGAA
AGCGCCTGCTGAGCTGGCGATCCGTCGATTGGCAAGAAAGCAGAAGGTGTTGGGCAAGGATTTCGACAAGGGGTCCGGA
AAACGACGGCAGCGGGCGGCAAGGGGACGTCAGCGACTCGACCAGGACCCCGCAAGGTCGGTGACGCAGGATGTCCGGG
TCTGGAGATACTGCTGGTCCAGGGCCGATGACGCCCCTGCTGACGGCCTCCCCGGCCGCTCACCAGCCTCCGAATTGAT
TTCGCGCTCAAGTCTTCGACAGGGAGCCCATTCATGCGCAAGGGCGAGGAGCTGTTCACCGGCGTCGTCCCGATCCTGG
TCGAGCTGGACGGCGACGTCAACGGCCACAAGTTCTCCGTCCGCGGCGAGGGCGAGGGCGACGCCACCAACGGCAAGCT
GACCCTGAAGTTCATCTGCACCACCGGCAAGCTGCCGGTCCCGTGGCCGACCCTGGTCACCACCCTGACCTACGGCGTC
CAGTGCTTCGCCCGCTACCCGGACCATATGAAGCAGCACGACTTCTTCAAGTCCGCCATGCCGGAGGGCTACGTCCAGG
AGCGCACCATCTCCTTCAAGGACGACGGCACCTACAAGACCCGCGCCGAGGTCAAGTTCGAGGGCGACACCCTGGTCAA
CCGCATCGAGCTGAAGGGCATCGACTTCAAGGAGGACGGCAACATCCTGGGCCACAAGCTGGAGTACAACTTCAACTCC
CACAACGTCTACATCACCGCCGACAAGCAGAAGAACGGCATCAAGGCCAACTTCAAGATCCGCCACAACGTCGAGGACG
GCTCCGTCCAGCTGGCCGACCACTACCAGCAGAACACCCCGATCGGCGACGGCCCGGTCCTGCTGCCGGACAACCACTA
CCTGTCCACCCAGTCCGTCCTGTCCAAGGACCCGAACGAGAAGCGCGACCACATGGTCCTGCTGGAGTTCGTCACCGCC
GCCGGCATCACCCACGGCATGGACGAGCTGTACAAGTGATGAACTAGTGTAGCTGACAAGAGACAGGATGAGGATCGTT
TCGCATGATTGAACAAGATGGATTGCACGCAGGTTCTCCGGCCGCTTGGGTGGAGAGGCTATTCGGCTATGACTGGGCA
CAACAGACAATCGGCTGCTCTGATGCCGCCGTGTTCCGGCTGTCAGCGCAGGGGCGCCCGGTTCTTTTTGTCAAGACCG
ACCTGTCCGGTGCCCTGAATGAACTGCAAGACGAGGCAGCGCGGCTATCGTGGCTGGCCACGACGGGCGTTCCTTGCGC
AGCTGTGCTCGACGTTGTCACTGAAGCGGGAAGGGACTGGCTGCTATTGGGCGAAGTGCCGGGGCAGGATCTCCTGTCA
TCTCACCTTGCTCCTGCCGAGAAAGTATCCATCATGGCTGATGCAATGCGGCGGCTGCATACGCTTGATCCGGCTACCT
GCCCATTCGACCACCAAGCGAAACATCGCATCGAGCGAGCACGTACTCGGATGGAAGCCGGTCTTGTCGATCAGGATGA
TCTGGACGAAGAGCATCAGGGGCTCGCGCCAGCCGAACTGTTCGCCAGGCTCAAGGCGCACATACCCGACGGCGAGGAT
CTCGTCGTGACCCATGGCGATGCCTGCTTGCCGAATATCATGGTGGAAAATGGCCGCTTTTCTGGATTCATCGACTGTG
GCCGGCTGGGTGTGGCGGACCGCTATCAGGACATAGCGTTGGCTACCCGTGATATTGCTGAAGAGCTTGGCGGCGAATG
GGCTGACCGCTTCCTCGTGCTTTACGGTATCGCCGCTCCCGATTCGCAGCGCATCGCCTTCTATCGCCTTCTTGACGAG
TTCTTCTGAGCGGGACTCTGGGGTAGCATAACCCCTTGGGGCCTCTAAACGGGTCTTGAGGGGTTTTTTGTTCGGGG

**Supplementary Figure 2. Organization of the reporter construct pSGK-choD for overproduction of cholesterol oxidase. a**, Plasmid map of pSGK-choD comprising two terminators (T4 Kurz and ECK120029600), a promoter (*P_choD*), a ribosome binding site (RBS), and two reporter genes (sf*gfp* and kanamycin) cloned into the pSET-152-based plasmid. **b**, Illustration of the construct along with the *choD* (*P_choD*) promoter cloned upstream of the reporter genes (sf*gfp* and kan). The two terminators (T4 Kurz and ECK) upstream of the *choD* promoter completely insulate the transcription leakage from the plasmid. *choD* promoter and its corresponding RBS drives the expression of sf*gfp* (super folder green fluorescent protein) and kanamycin resistance gene (*kan*). **c**, Depiction of nucleotide sequences of the region (*Xba*I/*Nde*I) highlighted in ‘b’ panel. Restriction enzymes (*Xba*I, *Nde*I, *Spe*I, and *Bam*HI) are highlighted with grey, T4 Kurz with plum color, ECK terminator with green; *choD* promoter with underlined in purple, sf*gfp* with bright green, kanamycin resistance gene with dark blue and bacteriophage T7-terminator with faint blue.

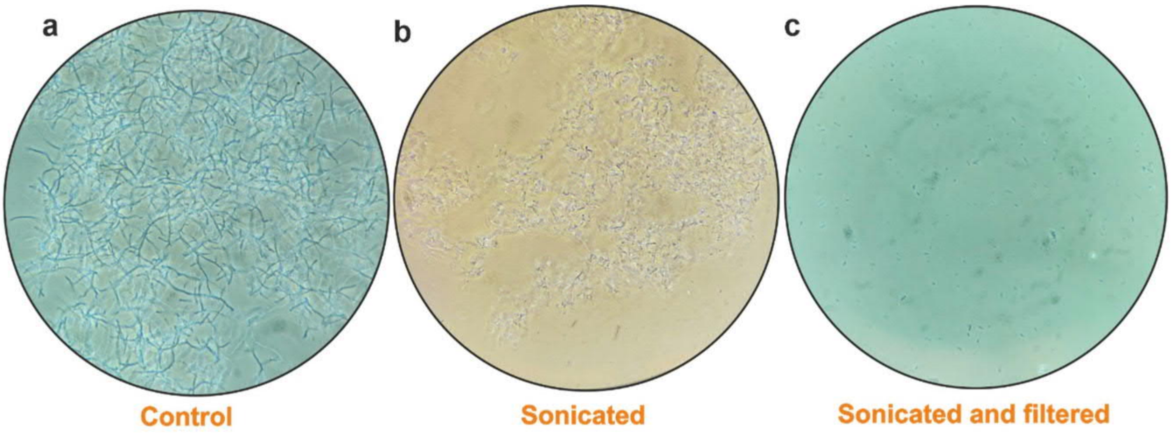

**Supplementary Figure 3. Example of fragmentation of *Amycolaptosis orientalis* mycelium.** **a**, *Amycolaptosis* cells growing in filamentous mycelium. **b**, Sonicated cells with mycelial filaments intertwined together. **c**, Sonicated samples filtered through cell-strainer cap falcon filter (Falcon® 12x75mm tube with cell strainer cap) display individual cells.

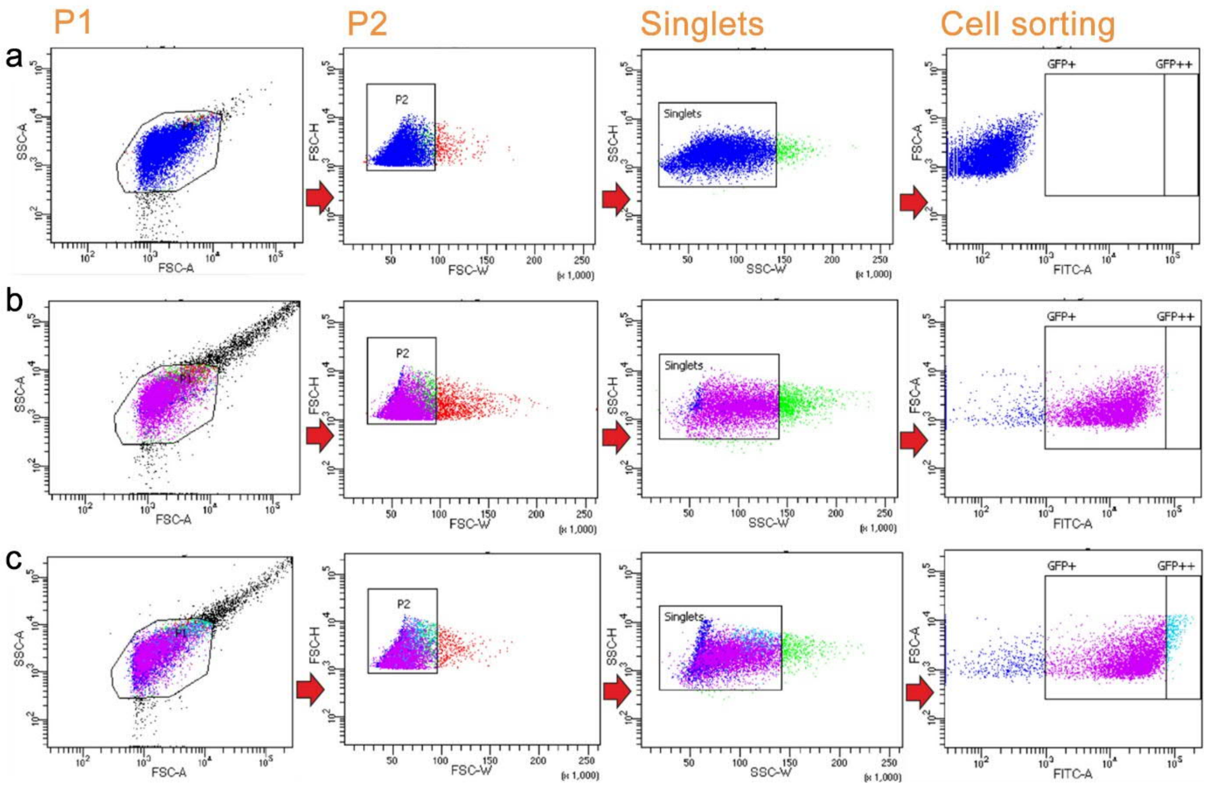

**Supplementary Figure 4. Example of FACS gating strategy and cell sorting of *Amycolaptosis orientalis*.** A two-step doublet discrimination gating was utilized to identify single cells. Initial single cell population, P1, was acquired through FSC-SSC dot plot gating. Signal heights and widths were taken into consideration in the second gating, P2, using FSC-H (heigth) and FSC-W (width) to identify a subpopulation of singlets. The singlets were sorted based on FITC-A fluorescence. **a**, Non-mutated *A. orientalis*/pSGKP45_UV0 was used as a negative control. **b**, *A. orientalis*/pS-GK was used as a positive control. **c**, Mutant library of *A. orientalis*/pSGKP45-UV1(3) were sorting to identify best *gfp* expressing clones. Each dot in the scatter plot represents a single cell. Legend: FITC, Fluorescence of singlets; FITC-A, Fluorescence area, FSC, Forward scatter; FSC-A, Forward scatter area; FSC-H, Forward scatter height; SSC, Side scatter; SSC-A, Side scatter area; SSC-H, Side scatter height, SSC-W, Side scatter width.

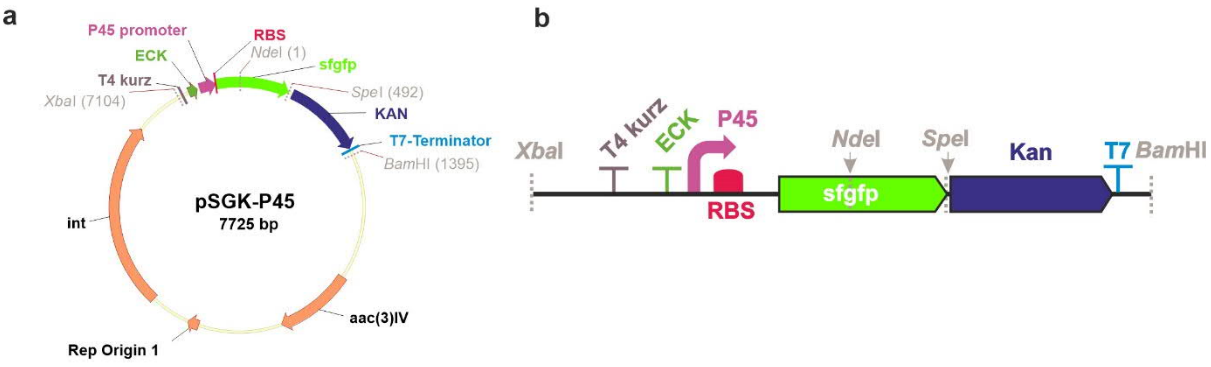

c

CTAGAGAGCTCGCTGAGTTGGCTGCTGCCACCGCTGAGCAATAACAAGCTTTATGCTTGTAAACCGTTTTGTGAAAAAA
TTTTTAAAATAAAAAAGGGGACCTCTAGGATTTTCAGCCAAAAAACTTAAGACCGCCGGTCTTGTCCACTACCTTGCAG
TAATGCGGTGGACAGGATCGGCGGTTTTCTTTTCTCTTCTCAACTGAAAGGAGGAACTATATCCGCTCCGTCCGCGCGT
ACGGCCAGCGGTCCCAATATTGCGATCTGCATCCGACCCCCAGATACCTGCCGAAAGAGCACGCCAGGCAGCTGCCCGC
GTGCTTCGCTTTCCCCCAGTGAGACGCCAGCGACGCTAGCAGAGAAACGGCTTCCTGATCCCGGGCGGGAAAAACATGC
GCAAGGGCGAGGAGCTGTTCACCGGCGTCGTCCCGATCCTGGTCGAGCTGGACGGCGACGTCAACGGCCACAAGTTCTC
CGTCCGCGGCGAGGGCGAGGGCGACGCCACCAACGGCAAGCTGACCCTGAAGTTCATCTGCACCACCGGCAAGCTGCCG
GTCCCGTGGCCGACCCTGGTCACCACCCTGACCTACGGCGTCCAGTGCTTCGCCCGCTACCCGGACCATATGAAGCAGC
ACGACTTCTTCAAGTCCGCCATGCCGGAGGGCTACGTCCAGGAGCGCACCATCTCCTTCAAGGACGACGGCACCTACAA
GACCCGCGCCGAGGTCAAGTTCGAGGGCGACACCCTGGTCAACCGCATCGAGCTGAAGGGCATCGACTTCAAGGAGGAC
GGCAACATCCTGGGCCACAAGCTGGAGTACAACTTCAACTCCCACAACGTCTACATCACCGCCGACAAGCAGAAGAACG
GCATCAAGGCCAACTTCAAGATCCGCCACAACGTCGAGGACGGCTCCGTCCAGCTGGCCGACCACTACCAGCAGAACAC
CCCGATCGGCGACGGCCCGGTCCTGCTGCCGGACAACCACTACCTGTCCACCCAGTCCGTCCTGTCCAAGGACCCGAAC
GAGAAGCGCGACCACATGGTCCTGCTGGAGTTCGTCACCGCCGCCGGCATCACCCACGGCATGGACGAGCTGTACAAGT
GATGAACTAGTGTAGCTGACAAGAGACAGGATGAGGATCGTTTCGCATGATTGAACAAGATGGATTGCACGCAGGTTCT
CCGGCCGCTTGGGTGGAGAGGCTATTCGGCTATGACTGGGCACAACAGACAATCGGCTGCTCTGATGCCGCCGTGTTCC
GGCTGTCAGCGCAGGGGCGCCCGGTTCTTTTTGTCAAGACCGACCTGTCCGGTGCCCTGAATGAACTGCAAGACGAGGC
AGCGCGGCTATCGTGGCTGGCCACGACGGGCGTTCCTTGCGCAGCTGTGCTCGACGTTGTCACTGAAGCGGGAAGGGAC
TGGCTGCTATTGGGCGAAGTGCCGGGGCAGGATCTCCTGTCATCTCACCTTGCTCCTGCCGAGAAAGTATCCATCATGG
CTGATGCAATGCGGCGGCTGCATACGCTTGATCCGGCTACCTGCCCATTCGACCACCAAGCGAAACATCGCATCGAGCG
AGCACGTACTCGGATGGAAGCCGGTCTTGTCGATCAGGATGATCTGGACGAAGAGCATCAGGGGCTCGCGCCAGCCGAA
CTGTTCGCCAGGCTCAAGGCGCACATACCCGACGGCGAGGATCTCGTCGTGACCCATGGCGATGCCTGCTTGCCGAATA
TCATGGTGGAAAATGGCCGCTTTTCTGGATTCATCGACTGTGGCCGGCTGGGTGTGGCGGACCGCTATCAGGACATAGC
GTTGGCTACCCGTGATATTGCTGAAGAGCTTGGCGGCGAATGGGCTGACCGCTTCCTCGTGCTTTACGGTATCGCCGCT
CCCGATTCGCAGCGCATCGCCTTCTATCGCCTTCTTGACGAGTTCTTCTGAGCGGGACTCTGGGGTAGCATAACCCCTT
GGGGCCTCTAAACGGGTCTTGAGGGGTTTTTTGTTCGGGG

**Supplementary Figure 5. Organization of the reporter construct pSGK-P45 for activation of mutaxanthene BGC. a**, Plasmid map of pSGK-P45 comprising two terminators (T4 Kurz and ECK (ECK120029600)), a *mutR* promoter (P45), a ribosome binding site (RBS), and two reporter genes (sf*gfp* and kanamycin) cloned into the pSET-152-based plasmid. **b**, Illustration of the construct along with the *mutR* promoter cloned upstream of the reporter genes (sf*gfp* and Kan). The two terminators (T4 Kurz and ECK) upstream of the *mutR* promoter completely insulate the transcription leakage from the plasmid. *mutR* promoter and its corresponding RBS drives the expression of sf*gfp* (super folder green fluorescent protein) and kanamycin resistance gene (*kan*). **c**, Depiction of nucleotide sequences of the region (*Xba*I/*Nde*I) highlighted in the 'b' panel. Restriction enzymes (*Xba*I, *Nde*I, *Spe*I, and *Bam*HI) are highlighted with grey, T4 Kurz with plum, ECK terminator with green; *mutR* promoter underlined in purple, sf*gfp* with bright green, kanamycin resistance gene with dark blue and bacteriophage T7-terminator with faint blue.

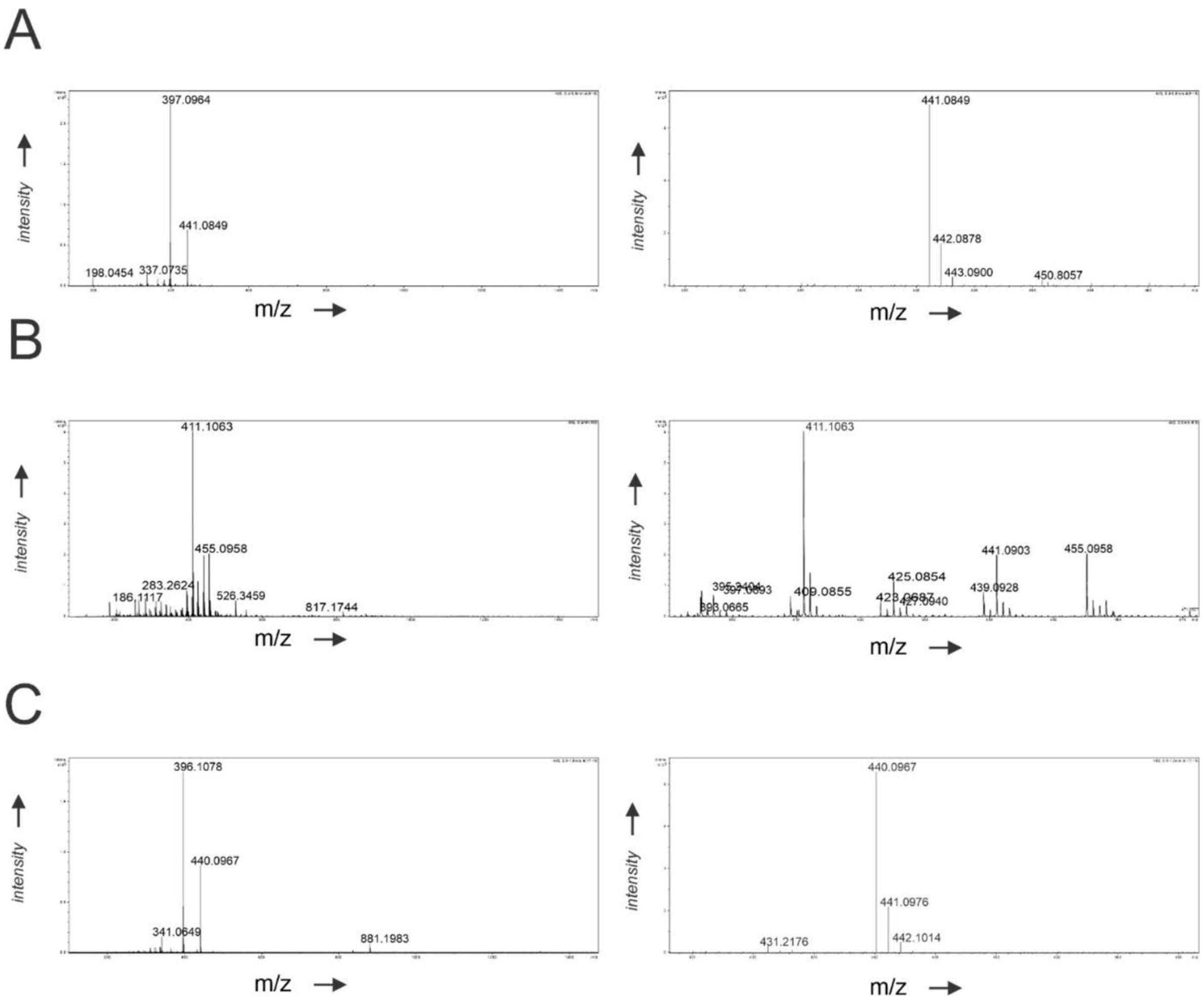

**Supplementary Figure 6. HR-MS spectra.** A) HR-MS spectrum of **1**. ESI m/z [M-H]- obs.441.0849, calc. 441.0827 B) HR-MS spectrum of **2**. ESI m/z [M-H]- obs. 455.0958, calc. 455.0984 C) HR-MS spectrum of **3**. ESI m/z [M-H]- obs. 440.0967, calc.440.0982.

**Supplementary Table 1.** Coding DNA Sequences from the silent gene cluster of *Streptomyces* sp. NRRL F3213 and their proposed roles within the biosynthetic pathway.

| Gene name | Size (AA) | RAST annotation | Most similar ORF [Species](Identity %) | Accession no. |
|---|---|---|---|---|
| ***mutA*** | 113 | Transcriptional regulator, PadR family | PadR family transcriptional regulator [*Nonomuraea* sp. WAC 01424](87) | WP_125636118.1 |
| ***mutB*** | 196 | Hypothetical protein | Hypothetical protein [*Kibdelosporangium aridum*](91) | WP_037276114.1 |
| ***mutC*** | 90 | Hypothetical protein | Predicted protein [*Streptomyces* sp. AA4] (97.78) | EFL10681.1 |
| ***mutD*** | 332 | Malonyl CoA-acyl carrier protein transacylase | Acyltransferase domain-containing protein [*Nonomuraea montanisoli*](54) | WP_175592186.1 |
| ***mutE*** | 344 | Hypothetical protein | Ketoacyl-ACP synthase III family protein [*Streptomyces* sp. Go-475](57) | WP_114255099.1 |
| ***mutF*** | 257 | Hypothetical protein | Cyclase family protein [*Kutzneria* sp. CA-103260](74) | WP_211768124.1 |
| ***mutG*** | 249 | Short-chain dehydrogenase/reductase SDR | SDR family oxidoreductase [*Nocardia arthritidis*](59.35) | QIS10038.1 |
| ***mutH*** | 617 | Long-chain-fatty-acid--CoA ligase | AMP-binding protein [*Kutzneria* sp. CA-103260](55) | WP_211768122.1 |
| ***mutI*** | 319 | Aromatase | Aromatase/cyclase [*Kutzneria* sp. CA-103260](69) | WP_211768121.1 |
| ***mutJ*** | 261 | Acetoacetyl-CoA reductase | SDR family NAD(P)-dependent oxidoreductase [*Kutzneria albida*](74) | WP_025361002.1 |
| ***mutK*** | 411 | Polyprenyl-6-methoxyphenol hydroxylase and related FAD-dependent oxidoreductases | FAD-dependent monooxygenase [*Kutzneria* sp. CA-103260](63) | WP_211768119.1 |
| ***mutL*** | 252 | 3-oxoacyl-[acyl-carrier protein] reductase | SDR family oxidoreductase [*Streptomyces thermodiastaticus*](52) | WP_189795586.1 |
| ***mutM*** | 188 | *O*-methyltransferase | Ubiquinone/menaquinone biosynthesis C-methylase UbiE [*Kutzneria viridogrisea*](47) | MBA8930871.1 |
| ***mutN*** | 509 | Putative integral membrane protein | EmrB/QacA subfamily drug resistance transporter [*Candidatus Frankia californiensis*](52) | SBW23174.1 |
| ***mutO*** | 85 | Acyl carrier protein | Acyl carrier protein [*Thermomonospora echinospora*](55) | WP_103936125.1 |
| ***mutP*** | 401 | Polyketide chain length factor WhiE-CLF paralog | Ketosynthase chain-length factor [*Streptomyces* sp. CBMA152](65) | WP_188275488.1 |
| ***mutQ*** | 430 | Polyketide beta-ketoacyl synthase WhiE-KS paralog | Beta-ketoacyl-[acyl-carrier-protein] synthase family protein [*Kutzneria albida*](75) | WP_030111238.1 |
| ***mutR*** | 680 | Regulatory protein | DNA-binding SARP family transcriptional activator [*Kutzneria viridogrisea*](46) | MBA8926443.1 |
| ***mutS*** | 251 | Oxidoreductase, short-chain dehydrogenase/reductase family | Ketoreductase [*Streptomyces argillaceus*](58) | CAK50776.1 |
| ***mutT*** | 516 | 2-polyprenyl-6-methoxyphenol hydroxylase and related FAD-dependent oxidoreductases | FAD-dependent oxidoreductase [*Kitasatospora* sp. NA04385](62) | WP_176190106.1 |
| ***mutU*** | 340 | *O*-methyltransferase | Methyltransferase [*Kutzneria albida*](68) | WP_025361012.1 |
| ***mutV*** | 499 | Rifampin monooxygenase | FAD-dependent monooxygenase [*Kutzneria* sp. CA-103260](64) | WP_211768107.1 |
| ***mutW*** | 1227 | Dihydrolipoamide succinyltransferase component (E2) of 2-oxoglutarate dehydrogenase complex | Alpha-ketoglutarate decarboxylase [*Amycolatopsis regifaucium*](87) | WP_081809737.1 |
| ***mutX*** | 149 | Hypothetical protein | Nitroreductase family deazaflavin-dependent oxidoreductase [*Nocardia miyunensis*](69) | WP_084531700.1 |
| ***mutY*** | 412 | Putative membrane protein | FAD-dependent oxidoreductase [*Pseudonocardia eucalypti*](67) | WP_185062231.1 |
| ***mutZ*** | 399 | Nitrate/nitrite transporter NarK | MFS transporter [*Prauserella sediminis*](81) | WP_183785025.1 |

**Supplementary Table 2.** $^{1}H$ and $^{13}C$ NMR data for **3** recorded in MeOD-$d_4$.

| position | δ ($^{1}$H, ppm) | δ ($^{13}$C, ppm) | *J* (Hz) |
|---|---|---|---|
| **1** | 3.91 | 86.4 | $J_{1,4a}$ = 0.5 |
| **2** | | 192.0 | |
| **3** | | 103.4 | |
| **4** | | 191.5 | |
| **4a** | 5.51 | 77.5 | $J_{4a,1}$ = 0.5, $J_{4a,12a}$ = 2.2 |
| **5** | | 155.2 | |
| **6** | | 184.6 | |
| **6a** | | 115.2 | |
| **7** | | 162.8 | |
| **8** | 7.22 | 124.8 | $J_{8,9}$ = 8.5, $J_{8,10}$ = 1.1 |
| **9** | 7.63 | 138.0 | $J_{9,8}$ = 8.5, $J_{9,10}$ = 7.4 |
| **10** | 7.52 | 119.8 | $J_{10,8}$ = 1.1, $J_{10.9}$ = 7.4 |
| **10a** | | 133.2 | |
| **11** | | 183.9 | |
| **11a** | | 120.2 | |
| **12** | a: 3.13; b: 2.38 | 23.3 | $J_{12a,12b}$ = –18.9, $J_{12a,4a}$ = 2.2 |
| **12a** | | 47.8 | |
| **13** | | 183.4 | |
| **14** | a: 2.91; b: 2.87 | 30.8 | $J_{14a,14b}$ = –12.9, $J_{14a,15}$ = $J_{14b,15}$ = 7.4 |
| **15** | 1.19 | 12.9 | $J_{15,14a}$ = $J_{15,14b}$ = 7.4 |
| **16** | | 173.4 | |
| **17** | 3.50 | 60.1 | |

**Supplementary Table 3.** $^{1}H$ and $^{13}C$ NMR data for **2** (major tautomer with C2 as enol and C4 as ketone) recorded in $CDCl_3$.

| **position** | **δ ($^{1}$H, ppm)** | **δ ($^{13}$C, ppm)** | ***J* (Hz)** |
|---|---|---|---|
| **1** | 4.15 | 81.2 | |
| **2** | | 189.7 | |
| **3** | | 107.6 | |
| **4** | | 187.1 | |
| **4a** | 5.54 | 75.8 | $J_{4a,12a}$ = 2.5 |
| **5** | | 154.4 | |
| **6** | | 183.3 | |
| **6a** | | 115.4 | |
| **7** | | 160.1 | |
| **8** | 7.18 | 117.7 (117.68) | $J_{8,9}$ = 8.4, $J_{8,10}$ = 1.1 |
| **9** | 7.56 | 135.2 | $J_{9,8}$ = 8.4, $J_{9,10}$ = 7.6 |
| **10** | 7.60 | 118.8 | $J_{10,8}$ = 1.1, $J_{10.9}$ = 7.6 |
| **10a** | | 133.8 | |
| **11** | | 176.5 | |
| **11a** | | 118.6 | |
| **12** | a: 3.19; b: 2.35 | 22.4 | $J_{12a,12b}$ = –18.7, $J_{12a,4a}$ = 2.5 |
| **12a** | | 47.1 | |
| **13** | | 207.8 | |
| **14** | a: 3.05; b: 2.96 | 33.7 | $J_{14a,14b}$ = –18.3, $J_{14a,15}$ = $J_{14b,15}$ = 7.2 |
| **15** | 1.12 | 8.1 | $J_{15,14a}$ = $J_{15,14b}$ = 7.2 |
| **16** | | 173.0 | |
| **17** | 3.50 | 60.9 | |
| **18** | 3.95 | 56.5 (56.47) | |

**Supplementary Table 4.** $^{1}$H and $^{13}$C NMR data for **2** (minor tautomer with C4 as enol and C2 as ketone) recorded in $CDCl_3$.

| **position** | **δ ($^{1}$H, ppm)** | **δ ($^{13}$C, ppm)** | ***J* (Hz)** |
|---|---|---|---|
| **1** | 3.97 | 84.1 | |
| **2** | | 192.8 | |
| **3** | | 108.5 | |
| **4** | | 187.9 | |
| **4a** | 5.77 | 72.3 | $J_{4a,12a}$ = 1.9 |
| **5** | | 154.0 | |
| **6** | | 183.2 | |
| **6a** | | 115.7 | |
| **7** | | 160.1 | |
| **8** | 7.18 | 117.7 (117.70) | $J_{8,9}$ = 8.5, $J_{8,10}$ = 1.2 |
| **9** | 7.56 | 135.3 | $J_{9,8}$ = 8.5, $J_{9,10}$ = 7.4 |
| **10** | 7.58 | 118.9 | $J_{10,8}$ = 1.2 $J_{10.9}$ = 7.4 |
| **10a** | | 133.7 | |
| **11** | | 176.4 | |
| **11a** | | 118.4 | |
| **12** | a: 3.16; b: 2.42 | 21.5 | $J_{12a,12b}$ = –18.9, $J_{12a,4a}$ = 1.9 |
| **12a** | | 46.8 | |
| **13** | | 207.1 | |
| **14** | a: 3.07 | 33.6 | $J_{14,15}$ = 7.2 |
| **15** | 1.15 | 8.2 | $J_{15,14}$ = 7.2 |
| **16** | | 173.4 | |
| **17** | 3.46 | 59.6 | |
| **18** | 3.95 | 56.5 (56.46) | |

**Supplementary Table 5.** $^1$H and $^{13}$C NMR data for **1** (major tautomer with C2 as enol and C4 as ketone) recorded in $CDCl_3$.

| position | δ ($^1$H, ppm) | δ ($^{13}$C, ppm) | *J* (Hz) |
|---|---|---|---|
| **1** | 4.18 | 81.3 | |
| **2** | | 189.5 | |
| **3** | | 107.5 | |
| **4** | | 187.0 | |
| **4a** | 5.58 | 75.6 | $J_{4a,12a}$ = 2.3 |
| **5** | | 153.5 | |
| **6** | | 182.8 | |
| **6a** | | 113.8 | |
| **7** | | 161.8 | |
| **8** | 7.11 | 124.2 | $J_{8,9}$ = 8.4, $J_{8,10}$ = 1.1 |
| **9** | 7.48 | 118.9 (118.93) | $J_{9,8}$ = 8.4, $J_{9,10}$ = 7.4 |
| **10** | 7.44 | 136.8 | $J_{10,8}$ = 1.1 $J_{10.9}$ = 7.4 |
| **10a** | | 131.4 | |
| **11** | | 182.4 | |
| **11a** | | 118.3 | |
| **12** | a: 3.24; b: 2.36 | 22.6 | $J_{12a,12b}$ = –19.0, $J_{12a,4a}$ = 2.3 |
| **12a** | | 47.1 | |
| **13** | | 207.9 | |
| **14** | a: 3.06; b: 2.98 | 33.7 | $J_{14a,14b}$ = –18.2, $J_{14a,15}$ = $J_{14b,15}$ = 7.2 |
| **15** | 1.13 | 8.1 | $J_{15,14a}$ = $J_{15,14b}$ = 7.2 |
| **16** | | 173.7 | |
| **17** | 3.62 | 60.8 | |

**Supplementary Table 6.** $^1$H and $^{13}$C NMR data for **1** (major tautomer with C4 as enol and C2 as ketone) recorded in $CDCl_3$.

| position | δ ($^1$H, ppm) | δ ($^{13}$C, ppm) | *J* (Hz) |
|---|---|---|---|
| **1** | 4.00 | 84.3 | |
| **2** | | 192.9 | |
| **3** | | 108.5 | |
| **4** | | 187.6 | |
| **4a** | 5.82 | 72.2 | $J_{4a,12a}$ = 2.0 |
| **5** | | 153.1 | |
| **6** | | 182.5 | |
| **6a** | | 113.6 | |
| **7** | | 161.7 | |
| **8** | 7.06 | 124.3 | $J_{8,9}$ = 8.4, $J_{8,10}$ = 1.1 |
| **9** | 7.41 | 136.8 | $J_{9,8}$ = 8.4, $J_{9,10}$ = 7.4 |
| **10** | 7.34 | 118.9 (118.89) | $J_{10,8}$ = 1.1 $J_{10.9}$ = 7.4 |
| **10a** | | 131.2 | |
| **11** | | 182.2 | |
| **11a** | | 118.6 | |
| **12** | a: 3.18; b: 2.39 | 21.6 | $J_{12a,12b}$ = –19.3, $J_{12a,4a}$ = 2.0 |
| **12a** | | 46.8 | |
| **13** | | 207.2 | |
| **14** | a: 3.10; b: 3.09 | 33.4 | $J_{14a,14b}$ = –17.9, $J_{14a,15}$ = $J_{14b,15}$ = 7.2 |
| **15** | 1.17 | 8.2 | $J_{15,14a}$ = $J_{15,14b}$ = 7.2 |
| **16** | | 174.4 | |
| **17** | 3.48 | 59.4 | |
